# Comparative genomics reveals modular virulence repertoires and extensive horizontal gene transfer in *Vibrio* species associated with white syndrome of *Porites cylindrica*

**DOI:** 10.64898/2026.03.19.713055

**Authors:** Ewelina Rubin, Gaurav Shimpi, Héloïse Rouzé, Bastian Bentlage

**Affiliations:** Marine Laboratory, University of Guam, Mangilao, GU 96923; Department of Ecosystem Studies, Graduate School of Agricultural and Life Sciences, The University of Tokyo, Yayoi, Tokyo, Japan; UMR MARBEC, University of Montpellier, CNRS, IFREMER, IRD, Mayotte

## Abstract

Coral white syndrome (WS) is a widespread condition characterized by tissue loss and skeletal exposure, with multiple bacterial pathogens—particularly *Vibrio* species—implicated in its etiology. To investigate the genomic basis of potential virulence in *Vibrio* associated with WS of *Porites cylindrica*, we isolated 57 *Vibrio* strains from healthy and diseased coral tissues collected from reefs in Guam. Thirteen representative strains from five dominant species—*V. coralliilyticus, V. owensii, V. tubiashii, V. harveyi,* and *V. tetraodonis*—were selected for whole-genome sequencing using Oxford Nanopore Technology. Comparative genomic analyses revealed a conserved repertoire of extracellular enzymes, including hemolysins, cytolysins, metalloproteases, and subtilisin-like peptidases, alongside lineage-specific toxin and regulatory modules. Variation in secretion systems (T1SS–T6SS), particularly in T3SS, T4SS, and T6SS subtypes, reflected diversification of host-interaction and competitive capacities across species. Mobile genetic elements, including plasmids and prophages, contributed additional virulence-associated genes and secretion clusters, underscoring the role of horizontal gene transfer in shaping the accessory genome content. Notably, *V. coralliilyticus* harbored cholera toxin–related genes (*ace* and *zot*), while conjugative plasmid systems indicated potential for gene dissemination across lineages. Together, these findings demonstrate that virulence potential in coral-associated *Vibrio* is broadly distributed and structured by a conserved ecological core overlaid with flexible, horizontally acquired modules. This study provides the first comparative genomic framework for *Vibrio* associated with *P. cylindrica*, advancing our understanding of how genomic plasticity and modular virulence repertoires may contribute to opportunistic disease dynamics in coral reef ecosystems.

## Introduction

*Porites cylindrica* is one of the dominant reef-building corals throughout the Indo-Pacific [1, 2]. This coral species is considered highly resilient, capable of recovering from both bleaching events and disease outbreaks [1, 3, 4]. In Guam, Micronesia, *P. cylindrica* accounts for up to 73% of coral cover across surveyed reefs [5, 6]. Despite its reputation for resilience, *P. cylindrica* is chronically affected by white syndrome (WS) lesions [7–9]. The causes of the WS lesions in *P. cylindrica* remain unknown. However, in other coral species, WS has been linked to multiple etiologies [10, 11]. Because of this, WS is not classified as a single disease but rather as a group of coral diseases that share a common primary symptom: tissue loss that exposes the white coral skeleton [12, 13]. This tissue loss associated with WS can be caused by environmental stress, pathogenic agents, or a combination of both, and can result in partial or complete mortality of coral colonies, both in the wild and in restoration projects [11, 13]. After tissue is lost, the bare white skeleton is often rapidly colonized by other organisms, primarily turf algae or cyanobacteria [14]. Further research is needed to elucidate the microbial communities involved, the mechanisms underlying *P. cylindrica* WS development and progression, and the causal pathways. With rising seawater temperatures and increasing anthropogenic pressure, coral disease outbreaks are expected to become more frequent and severe [15]. Understanding the causes of coral disease events and their impacts is crucial for developing effective mitigation strategies, which require molecular tools for diagnostics and monitoring, as well as advances in treatments and interventions.

Bacterial pathogens are commonly investigated as possible drivers of coral white syndromes. For instance, WS in *Pocillopora damicornis* can be triggered by opportunistic infection with *Vibrio coralliilyticus* strain BAA-450 during periods of immune suppression caused by elevated seawater temperatures [10]. However, WS lesions in *P. damicornis* can also result from infection with *V. harveyi* [16], demonstrating the multi-etiology of WS in the same host. Similarly, six *Vibrio* isolates have been identified as the primary pathogens responsible for WS in three different coral species (*Pachyseris speciosa, Montipora aequituberculata*, and *Acropora cytherea*) across the Indo-Pacific [17]. One of these isolates was later identified through genome sequencing as *V. coralliilyticus* strain P1 [18]. Other pathogenic *Vibrio* strains include *V. coralliilyticus* strain OCN008, which causes acute WS in *Montipora capitata* [19], and *V. owensii* strain OCN002, which causes a chronic form of WS in the same host [20]. *V. tubiashii* has been implicated in WS in *Porites lutea* [14], while *V. alginolyticus* is associated with WS in *Porites andrewsi* [21]. These examples demonstrate that various *Vibrio* species are associated with WS in multiple coral hosts. In addition, the microbiome study of *P. cylindrica* lesions shows that the genus *Vibrio* was detected in 58% of WS lesions in corals from two reef sites in Guam [22], however nothing is known about the genomic diversity of *Vibrio* lineages associated with these lesions.

The genus *Vibrio* is large and highly diverse, with close to 150 described species that are abundant in estuarine and marine environments. In addition to being pathogenic to corals, many *Vibrio* species (e.g., *V. cholerae, V. vulnificus*, and *V. parahaemolyticus*) are pathogenic to humans and to aquatic animals, such as fish (e.g., *V. harveyi*, *V. alginolyticus*, and *V. ponticus*), crustaceans (e.g., *V. owensii*), mollusks (e.g., *V. coralliilyticus*) [23–30]. Given their clinical and economic impact, the molecular mechanisms underlying *Vibrio* pathogenicity have been extensively studied [31–34]. Many *Vibrio* strains possess a variety of virulence factors (VFs) defined as molecules that allow them to identify, invade, colonize, and spread within a host. However, not all strains of a *Vibrio* species are pathogenic; frequently, pathogenicity is acquired through horizontal gene transfer (HGT). For example, pathogenic *V. cholerae* strains that cause cholera acquired the cholera toxin (CT) gene via HGT mediated by a bacteriophage [24]. Advances in genome sequencing and comparative genomics have greatly improved our understanding of *Vibrio* pathogenicity and the role of HGT. Comparisons between pathogenic and non-pathogenic strains or species provide valuable insights into infection strategies, ecological adaptations, and unique genetic features. For instance, comparative genomic analysis of two *V. coralliilyticus* strains, P1 and BAA-450, revealed distinct pathogenicity islands and virulence factors [35]. Another study comparing four *V. coralliilyticus* strains pathogenic to both corals and oyster larvae showed that prophage-mediated HGT contributes to the spread of virulence within the *Vibrio* population [36]. A broader comparative analysis of more than 20 *Vibrio* strains from various species revealed that chitin metabolism is a core feature of the genus, with some species exhibiting expanded chitinase gene families, reflecting their adaptation to chitin-rich environments [37]. Similarly, analysis of a virulent *Vibrio* strain causing acute hepatopancreatic necrosis disease in shrimp identified a plasmid containing two genes encoding a deadly toxin [38]. The select examples above demonstrate the impact of comparative genomics and its importance in understanding *Vibrio* pathogenicity.

Despite the frequent association of *Vibrio* species with coral WS across multiple coral hosts, the genomic diversity and modular virulence architecture of *Vibrio* strains associated with WS lesions in *P. cylindrica* remain poorly resolved. In particular, it is unclear how virulence-associated genes, secretion systems, and horizontally acquired elements are distributed across strains isolated from healthy and diseased tissues, and whether pathogenic potential is confined to specific lineages or broadly embedded within a flexible accessory gene pool. Here, we use comparative genomics to analyze representative strains from five dominant *Vibrio* species isolated from both healthy and WS-affected tissues of *P. cylindrica* collected from Guam reefs. By characterizing their virulence repertoires, secretion system architectures, and mobile genetic elements, we aim to identify conserved and lineage-specific modules that may shape host interaction strategies. This approach provides a genomic framework for evaluating the role of *Vibrio* in WS pathogenesis and for understanding how horizontal gene transfer and genomic plasticity contribute to coral disease dynamics under changing environmental conditions.

## Materials and Methods

### Bacterial isolation

Seventeen coral samples, each approximately 2 cm in length, were collected from ten colonies of *P. cylindrica* at two shallow (1m – 2m depth) reef flat sites in Guam: Tumon Bay and Luminao. At Tumon Bay, all four sampled colonies displayed white syndrome (WS) lesions. From each of these colonies, one sample was taken from healthy tissue and one from diseased tissue. In contrast, at Luminao, three of the six sampled colonies exhibited WS lesions, while the other three appeared healthy. For the colonies with WS, both healthy and diseased tissues were collected, whereas only healthy tissues were sampled from the disease-free colonies. Coral branches were cut using sterilized wire cutters, which were cleaned between samples by immersion in 10% bleach followed by 95% ethanol. The samples were placed in sterile plastic bags and kept on ice during transport to the Marine Laboratory at the University of Guam. In the lab, each coral branch was rinsed with 0.2-µm-filtered, sterile water to limit seawater-associated surface bacteria. The coral tissue was then scraped off using sterilized scalpels and tweezers. The resulting tissue slurry was transferred into sterile microcentrifuge tubes and streaked onto culture plates for bacterial culture. To maximize the diversity of cultivated bacteria, four different types of culture media were used: (1) marine agar (MA), (2) peptone seawater media (PEP), (3) tryptone seawater media (TRY), and (4) thiosulfate citrate bile salts agar (TCBS). MA was prepared using Marine Broth 2216 (Millipore) and agar (Wako) according to the manufacturers’ protocols. TCBS (BD) was prepared based on the manufacturer’s protocol. The PEP medium consisted of 3 g of BactoTM proteose peptone (BD), 2 g of BactoTM yeast extract (Difco), and 15 g of agar (Wako) per liter of 0.2 µm-filtered, sterilized seawater. The TRY medium consisted of 3 g BactoTM tryptone (BD), 2 g BactoTM yeast extract (Difco), and 15 g agar (Wako) per liter of 0.2 µm-filtered, sterilized seawater. Each coral tissue slurry was streaked in duplicate on all four media types. From each plate, between three and five single bacterial colonies were picked and subcultured onto media plates at least three times to ensure the isolation of pure, axenic cultures. Liquid cultures of the isolated bacteria were prepared in the same medium without agar; they were preserved as glycerol stocks at a final glycerol concentration of 15% and stored at –80°C.

### Bacterial identification

Liquid cultures, each 3 mL, were grown for 12 to 18 hours at 28 °C. Following incubation, bacterial cells were pelleted by centrifugation, and DNA was extracted from the pellets using the GenCatch Blood & Tissue Genomic Mini Prep Kit (Epoch Life Science). The 16S rRNA gene was amplified using the universal bacterial primers 27F and 1492R under the PCR cycling conditions described by Galkiewicz and Kellogg (2008) [39]. PCR amplifications were performed using GoTaq Flexi Taq Polymerase and reagents according to the manufacturer’s protocol for a 25 ul reaction (Promega, Madison, WI). The PCR cycling conditions were as follows: an initial denaturation step for 2 min at 95 °C, followed by 34 cycles of 95 °C for 45 s, 50 °C for 45 s, and 72 °C for 90 s, with a final extension of 5 min at 72 °C. For isolates identified as *Vibrio*, a second PCR amplification targeting the RNA polymerase gene (*rpoA*) was performed to provide higher-resolution species-level identification. This second PCR was performed using the primers and PCR conditions described by Thompson et al. (2005) [40]. PCR amplifications for the *rpoA* gene were performed using GoTaq® Flexi Taq Polymerase and reagents following the manufacturer’s protocol for a 25 ul reaction (Promega, Madison, WI), and the PCR cycling conditions were as follows: an initial denaturation step for 2 min at 95 °C, followed by 30 cycles of 95 °C for 30 s, 50 °C for 30 s, and 72 °C for 60 s, with a final extension of 5 min at 72 °C. The resulting PCR products were purified using the GeneJET PCR Purification Kit (Thermo Fisher, Vilnius, Lithuania) and sent to Epoch Life Science Inc. services (Missouri City, Texas) for Sanger sequencing. The obtained sequences were trimmed for quality control using Geneious software, and genus-level taxonomic identification was performed using 16S sequences in BLAST searches against the 16S rRNA Silva database v. 138.2 [41], with 100% pairwise nucleotide identity and 100% query coverage. For *Vibrio* species, the *rpoA* gene was used in a BLAST search against the NCBI nt core nucleotide database, and species identification was determined using 100% query coverage and ≥99.5% nucleotide identity.

The final species-level identifications of 13 strains selected for genome sequencing were confirmed using multilocus sequence analysis (MLSA) following Thompson et al. (2005) [40] in combination with average nucleotide identity (ANI) values calculated using FastANI [42]. For phylogenetic analysis, eight housekeeping genes (*ftsZ*, *gap*, *gyrB*, *pyrH*, *recA*, *topA*, *rpoA*, and *mreB*) were extracted from the 13 annotated genomes generated in this study and 176 reference genomes downloaded from NCBI (Supplementary File 1). Individual gene alignments were constructed using MAFFT [43] implemented in Geneious and concatenated into a single supermatrix. Maximum-likelihood phylogenetic inference was performed using W-IQ-TREE [44, 45] with automatic model selection (ModelFinder). Branch support was assessed using 1,000 ultrafast bootstrap replicates.

### Whole genome sequencing

A total of thirteen *Vibrio* strains were selected for whole-genome sequencing. These strains represented the most frequently isolated *Vibrio* species and, when possible, included isolates from both reef sites and healthy and diseased coral tissues. Glycerol stocks of the selected strains were cultured in the original isolation media. Bacterial cells were then pelleted by centrifugation, and high-molecular-weight DNA was extracted using the Monarch HMW DNA Extraction Kit (New England Biolabs, Ipswich, MA). Sequencing libraries were prepared using the Ligation Sequencing Kit and the Native Barcoding Kit 24 from Oxford Nanopore Technologies (Oxfordshire, UK). Sequencing was performed on an R10 Nanopore flow cell (FLO-MIN114) paired with kit 14 chemistry on a MinION device over a 48-hour run time. The raw signal data (fast5 files) were base-called using Guppy basecaller version 2.2.3 (Oxford Nanopore Technologies) with the super-accuracy base-calling model in MinKNOW (version: 24.02.16.).

### Genome assembly and annotation

Genome assemblies were generated using Flye version 2.9.5 with the flag- -nano-hq and all the rest set to default settings [46], and the completeness of the assemblies was evaluated using BUSCO [47], with the reference lineage set to vibrio_odb12. Genome annotations were obtained using the Rapid Annotation using Subsystem Technology (RAST) server [48]. The annotated protein-coding sequences, provided in FASTA format by the RAST server, were analyzed using VFanalyzer [49] to identify virulence factor homologs shared with other pathogenic *Vibrio* species. These protein sequences were also submitted to the BlastKOALA server [50] to obtain KEGG identifiers for functional annotation. Protein sequences annotated by RAST as peptidases or proteases were further analyzed by comparing them with the MEROPS peptidase database [51], using local BLAST searches. Peptidases were assigned to the MEROPS family with a cutoff > 40% amino acid identity using a minimum of 70% query coverage. To predict horizontally transferred genomic regions, IslandViewer4 [52] and AlienHunter [53] were employed. The latter tool was run on the Proksee genome visualization platform [54]. To evaluate predicted horizontally transferred (HGT) regions, whole-chromosome sequence comparisons were performed using the BLAST ring alignment functionality implemented in the Proksee genome visualization platform [54]. For each chromosome and plasmid, the assembled genome was used as a reference sequence and compared with closely related *Vibrio* genomes retrieved from NCBI, using Average Nucleotide Identity (ANI >95% when available). BLASTN alignments were generated using default Proksee parameters, and regions lacking significant nucleotide similarity to reference genomes were identified as candidate strain-specific insertions. Bacteriophage sequences were identified using the online PHASTER tool [55]. To assess plasmid-mediated HGT, plasmid sequences were queried against the NCBI database using BLAST to identify homologous plasmids. We selected plasmid hits that met the following criteria (≥85% nucleotide identity and> 20% query coverage) and used these plasmids to conduct Average Nucleotide Identity (ANI) analysis. Finally, plasmids with ANI>85% were aligned and visualized using the Prokseek tool. Secretion system components identified in the *Vibrio* genomes were grouped according to the KEGG BRITE classification system (available at https://www.kegg.jp/brite/ko02044). This system enabled us to classify secretion-associated proteins into six major secretion systems and their respective subtypes. Specifically, the Type II Secretion System (T2SS) was further divided into four subsystems: the general secretion pathway (T2SS-GSP), the secretion and assembly of mannose-sensitive hemagglutinin pili (T2SS-MSHA), the secretion and assembly of other pili or fimbriae (T2SS-PILIN), and the secretion and assembly of tight adherence pili (T2SS-TAD). Similarly, the Type III Secretion System was subdivided into components related to flagellar assembly (T3SS-FLA) and those related to secretion systems distinct from flagella (T3SS). The Type IV Secretion System was divided into three categories: two associated with the assembly of conjugative pili required for plasmid transfer (T4SS-CON-P and T4SS-TRB), and one related to the transfer of DNA and proteins not associated with plasmid DNA (T4SS-DNA-PR). Figures showing gene arrangements in secretion systems were constructed using the Proksee tool or the gggenes package [56] in R version 4.5.1.

## Results

### *Vibrio* isolates and genome characteristics

A total of 115 bacterial strains were isolated from both healthy and diseased tissues of *P. cylindrica*, including representatives of Gammaproteobacteria, Alphaproteobacteria, Bacteroidia, and Actinobacteria (Supplementary Table S1). Among these, 57 isolates (49%) were identified as *Vibrio*. The most frequently isolated *Vibrio* species were *V. owensii* (11 isolates), *V. tubiashii* (11 isolates), members of the *V. aquimaris/tetraodonis* clade (11 isolates with 99% nucleotide identity of the *rpoA* gene between the two species), *V. coralliilyticus* (10 isolates), and *V. harveyi* (5 isolates) (Table 1). *V. tubiashii* was recovered exclusively from corals originating from Tumon (Table 1).

**Table 1.** Number of *Vibrio* strains per species isolated from healthy (H) and diseased (D) tissue of *P. cylindrica* colonies originating from two reef sites: Tumon (TU) and Luminao (LU)

|  | Number of isolates | TU-D | LU-D | TU-H | LU-H |
| --- | --- | --- | --- | --- | --- |
| *Vibrio owensii* | 11 | 2 | 4 | 4 | 1 |
| *Vibrio tubiashii* | 11 | 10 | 0 | 1 | 0 |
| *Vibrio tetraodonis/aquimaris* | 11 | 0 | 10 | 1 | 0 |
| *Vibrio coralliilyticus* | 10 | 7 | 2 | 0 | 1 |
| *Vibrio harveyi* | 5 | 4 | 0 | 1 | 0 |
| *Vibrio maritimus* | 4 | 1 | 3 | 0 | 0 |
| *Vibrio ponticus* | 1 | 0 | 0 | 0 | 1 |
| *Vibrio ishigakensis* | 1 | 0 | 0 | 0 | 1 |
| *Vibrio neptunius* | 1 | 1 | 0 | 0 | 0 |
| *Vibrio* spp. | 2 | 0 | 1 | 1 | 0 |
| total | 57 | 25 | 20 | 8 | 4 |

Of the 57 *Vibrio* isolates, 41 were obtained from diseased coral tissue and 16 from visually healthy tissue. Representative strains from the five most frequently isolated species were selected for whole-genome sequencing. The genome assemblies of the 13 sequenced strains were highly complete, with BUSCO scores exceeding 99% and sequencing coverage ranging from 52× to 148× (Table 2). Species assignments were confirmed using an eight-gene phylogenetic analysis (Supplementary Figure S1, Supplementary File 1) and average nucleotide identity (ANI >95%) (Supplementary Table S2). The isolates LU3DMA2 and LU5DMA4 were most closely related to the type strain of *V. tetraodonis* (95.19–95.47% ANI), whereas ANI values relative to *V. aquimaris* were below 95% (Supplementary Table S2). Twelve of the thirteen sequenced strains exhibited the typical bipartite *Vibrio* genome organization consisting of two chromosomes. The larger chromosomes ranged from 3.26 to 3.80 Mbp, and the smaller chromosomes ranged from 1.27 to 2.47 Mbp (Table 2). In contrast, strain LU5DMA4 (*V. tetraodonis*) possessed a single chromosome of approximately 4.76 Mbp (Table 2). Six strains harbored plasmids. Two strains each of *V. coralliilyticus*, *V. owensii*, and *V. tubiashii* carried a single plasmid, and one *V. tubiashii* strain contained two plasmids (Table 2). The remaining strains lacked plasmids.

**Table 2:** Genome characteristics of thirteen *Vibrio* strains belonging to five species isolated in this study. The strain label indicates: 1) location (LU or TU), 2) sample number, 3) tissue type (e.g., 5D – sample number 5 diseased tissue, 4H –sample number 4 healthy tissue), 4) isolation media (e.g., MA -marine agar, PEP – peptone, TRY- tryptone)

| Species | Strain | Tissue Type | Site | Genome length | Mean coverage | No. chromosomes | No. plasmids | BUSCO score |
| --- | --- | --- | --- | --- | --- | --- | --- | --- |
| *Vibrio tetraodonis* | LU5DMA4 | D | LU | 4,761,502 | 78 | 1 | 0 | 98.2% |
|  | LU3DMA2 | D | LU | 4,542,275 | 92 | 2 | 0 | 98.0% |
| *Vibrio coralliilyticus* | TU5DMA1 | D | TU | 5,796,936 | 102 | 2 | 1 | 99.5% |
|  | LU3DPEP4 | D | LU | 5,777,417 | 148 | 2 | 1 | 99.7% |
| *Vibrio harveyi* | TU3DPEP3 | D | TU | 5,798,052 | 52 | 2 | 0 | 99.1% |
|  | TU4HPEP4 | H | TU | 5,805,957 | 138 | 2 | 0 | 99.2% |
| *Vibrio owensii* | TU8HTRY5 | H | TU | 5,977,422 | 118 | 2 | 0 | 99.5% |
|  | TU3DMA2 | D | TU | 6,461,482 | 87 | 2 | 1 | 99.2% |
|  | LU5DTRY1 | D | LU | 6,100,886 | 121 | 2 | 1 | 99.2% |
|  | LU8HTRY1 | H | LU | 5,943,102 | 89 | 2 | 0 | 98.1% |
| *Vibrio tubiashii* | TU3DMA1 | D | TU | 5,684,442 | 104 | 2 | 2 | 99.5% |
|  | TU1DMA1 | D | TU | 5,318,319 | 145 | 2 | 1 | 99.7% |
|  | TU1DMA2 | D | TU | 5,423,814 | 162 | 2 | 1 | 99.6% |

### Virulence-associated genes

Comparative genomic analyses were performed to assess the distribution of previously characterized *Vibrio* virulence-associated genes across thirteen strains isolated from healthy and diseased *P. cylindrica* tissues. In total, 17 distinct virulence-associated gene categories were identified (Figure 1). These genes were classified as (i) lineage-restricted or strain-specific, or (ii) broadly distributed across most or all strains,

**Figure 1.**
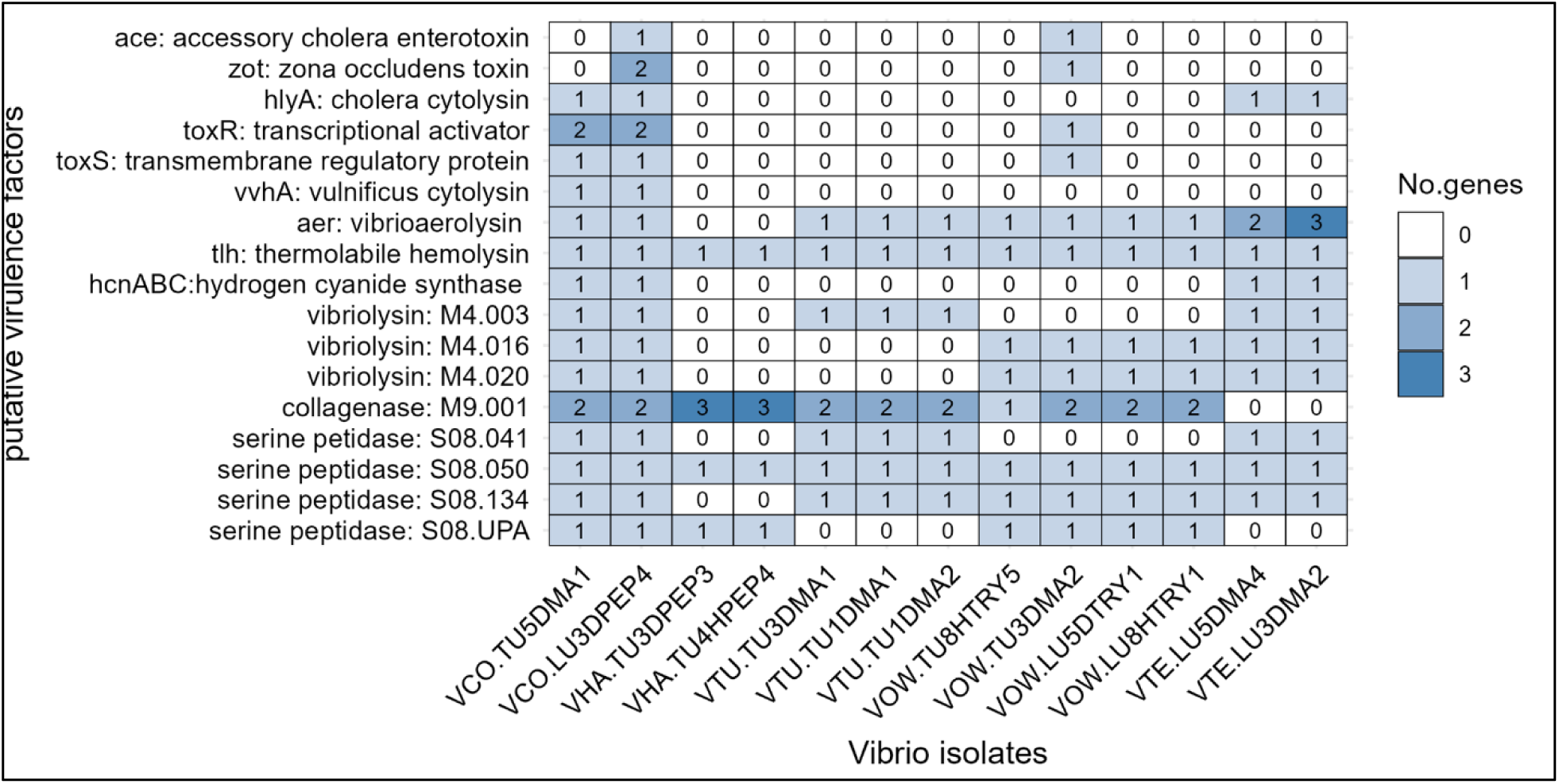
Distribution and copy number of predicted virulence-associated genes across thirteen *Vibrio* strains isolated from healthy and white syndrome (WS) lesion tissues of *Porites cylindrica*. Heatmap values indicate the number of predicted homologs per genome (0–3 copies). Strains are grouped by species: *Vibrio coralliilyticus* (VCO), *V. harveyi* (VHA), *V. tubiashii* (VTU), *V. owensii* (VOW), and *V. tetraodonis* (VTE). Putative virulence-associated genes were identified using the Virulence Factors Database (VFDB) and VFanalyzer, which predicts orthologous relationships to curated virulence factors based on sequence similarity and genomic context. Absence (0) indicates no detectable homolog under the applied search criteria.

#### Lineage-restricted and strain-specific virulence genes

Homologs of the *V. cholerae* cytolysin *hlyA* were identified in *V. coralliilyticus* and *V. tetraodonis*. These homologs shared 40–49% amino acid identity with *V. cholerae hlyA* but lacked the beta-prism lectin domain (Figure S3A). A homolog of the *V. vulnificus* hemolysin vvhA was detected exclusively in *V. coralliilyticus* strains, and those showed 76% amino acid identity to *V. vulnificus* (Figure 1; Figure S2B). Other cholera-associated genes, including *ace*, *zot*, *toxR*, and *toxS*, were also detected (Figure 1; Figure S3). The *ace,* homolog in strain VCO.LU3DPEP4 showed 57% amino acid identity to *V. cholerae* KWW43308, whereas the *ace*-like gene in strain VOW.TU3DMA2 showed 25% amino acid identity (Figure S3C). Two genes containing the zonular occludent domain (*zot*) were present in the *V. coralliilyticus* strain LU3DPEP4 and showed 53% amino acid identity to *V. cholerae* protein WP_000021609, and 68% identity to WP_275082003 (Figure 1, Figure S3D). The *hcnABC* gene cluster, encoding hydrogen cyanide synthases, was identified only in *V. coralliilyticus* and *V. tetraodonis* strains, with protein identities ranging from 42 to 61% to the hcnABC proteins of *Pseudomonas aeruginosa* PAO1 (Figure 1, Figure S3E).

#### Common and broadly distributed virulence factors

Several extracellular proteases were broadly distributed across multiple taxa (Figure 1). These included vibrio-aerolysins, metalloproteases (vibriolysins), collagenases, and subtilisins. M4.003-type vibriolysins were detected in *V. coralliilyticus*, *V. tubiashii*, and *V. tetraodonis*, whereas related M4.016 and M4.020 subtypes were identified in *V. coralliilyticus*, *V. tetraodonis*, and *V. owensii* (Figure 1). Collagenases (M9.001 subfamily) were present in most species, except *V. tetraodonis* and *V. harveyi,* which harbored three distinct collagenase genes (Figure 1). Subtilisin-type serine proteases (MEROPS S8 family) were identified in all strains of except strains of *V. harveyiii*, including S08.134-like homologs of *vvp1*/vvsa/asp (Figure 1). Additional subtilase subtypes (S08.41, S08.50, S08.UPA) were also detected and predicted to contain signal peptides indicating secretion (Figure 1). S08.50-like subtilases were present in all strains and showed similarity to *vhp1* from *V. harveyi*. The phospholipase-type cytolysin (*tlh)* was universally present across all thirteen strains (Figure 1).

### Distribution and diversity of secretion systems

Comparative genomic analysis of 13 coral-associated *Vibrio* strains identified multiple secretion systems typical of Gram-negative bacteria, including T1SS-T6SS (Figures 2–4). All strains encoded gene clusters required for flagellar assembly (T3SS-FLA). However, the number and organization of clusters varied among species (Figure 2). *V. harveyi* and *V. owensii* strains possessed two flagellar gene clusters: one encoding polar flagella (approximately 32 genes) on the large chromosome and a second encoding lateral flagella (26–29 genes) on the smaller chromosome (Figure 2). All three *V. tubiashii* strains contained two gene clusters (27+4 genes), both located on the large chromosome. The two *V. coralliilyticus* strains differed in the number of clusters (three and four, respectively). In addition to polar flagellar clusters (26+4 genes), both strains contained 41-gene clusters consistent with lateral flagellar systems. One *V. coralliilyticus* strain (LU3DPEP4) harbored an additional 23-gene flagellar assembly cluster within a predicted HGT region; based on gene count, this cluster appears incomplete. All strains encoded conserved T2SS-MSHA and T2SS-PILIN systems (Figure 2). In contrast, T2SS-TAD systems showed substantial variability. A total of 36 T2SS-TAD gene clusters were identified across 13 genomes, ranging from one to five per strain (Figure 2B). The highest copy number (five TAD clusters) was observed in two *V. tubiashii* strains, whereas one *V. owensii* strain contained a single TAD cluster (Figure 2B). Canonical T3SS injectosome clusters were identified in six strains belonging to *V. coralliilyticus*, *V. tetraodonis*, and *V. harveyi* (Figures 2 and 3). The T3SS clusters in *V. coralliilyticus* and *V. tetraodonis* resembled T3SS3, containing homologs of SseBCD translocation proteins and Salmonella-associated effectors (Figure 3, classification after Zakaria et al., 2023 [57]). In contrast, *V. harveyi* strains harbored T3SS clusters similar to Yersinia-like T3SS1 systems (Figure 3). Multiple T4SS subtypes were identified, frequently associated with plasmids (Figure 2). The T4SS-CON-P system was detected on plasmids in four strains and encoded genes consistent with conjugative pilus formation. A T4SS-TRB operon was identified on plasmid pVOW-LU5DTRY1-63 in *V. owensii*. Additionally, a T4SS-DNA-PR subtype was identified in four strains (two *V. harveyi* and two *V. owensii*). All strains encoded T6SS clusters, although the number of subtypes varied by lineage (Figure 4). *V. owensii* and *V. harveyi* strains possessed three T6SS subtypes, whereas *V. coralliilyticus*, *V. tetraodonis*, and *V. tubiashii* strains harbored two. Four structural T6SS subtypes (named after the first and last gene) were identified: 1) TssM–VgrG (present in all strains), 2) TssL–VasH (present in 11 strains), 3) PpkA–PAAR (restricted to *V. owensii* and *V. harveyi*), 4) VasL–PAAR (restricted to two *V. tubiashii* strains), (Figure 4). The TssM–VgrG subtype was typically located on chromosome 1, while TssL–VasH was located on chromosome 2. Searches for nine previously described anti-eukaryotic T6SS2 effectors identified eight of nine effectors in *V. coralliilyticus* strains. Homologs of several effectors were also detected in *V. tubiashii* and *V. tetraodonis* strains.

**Figure 2.**
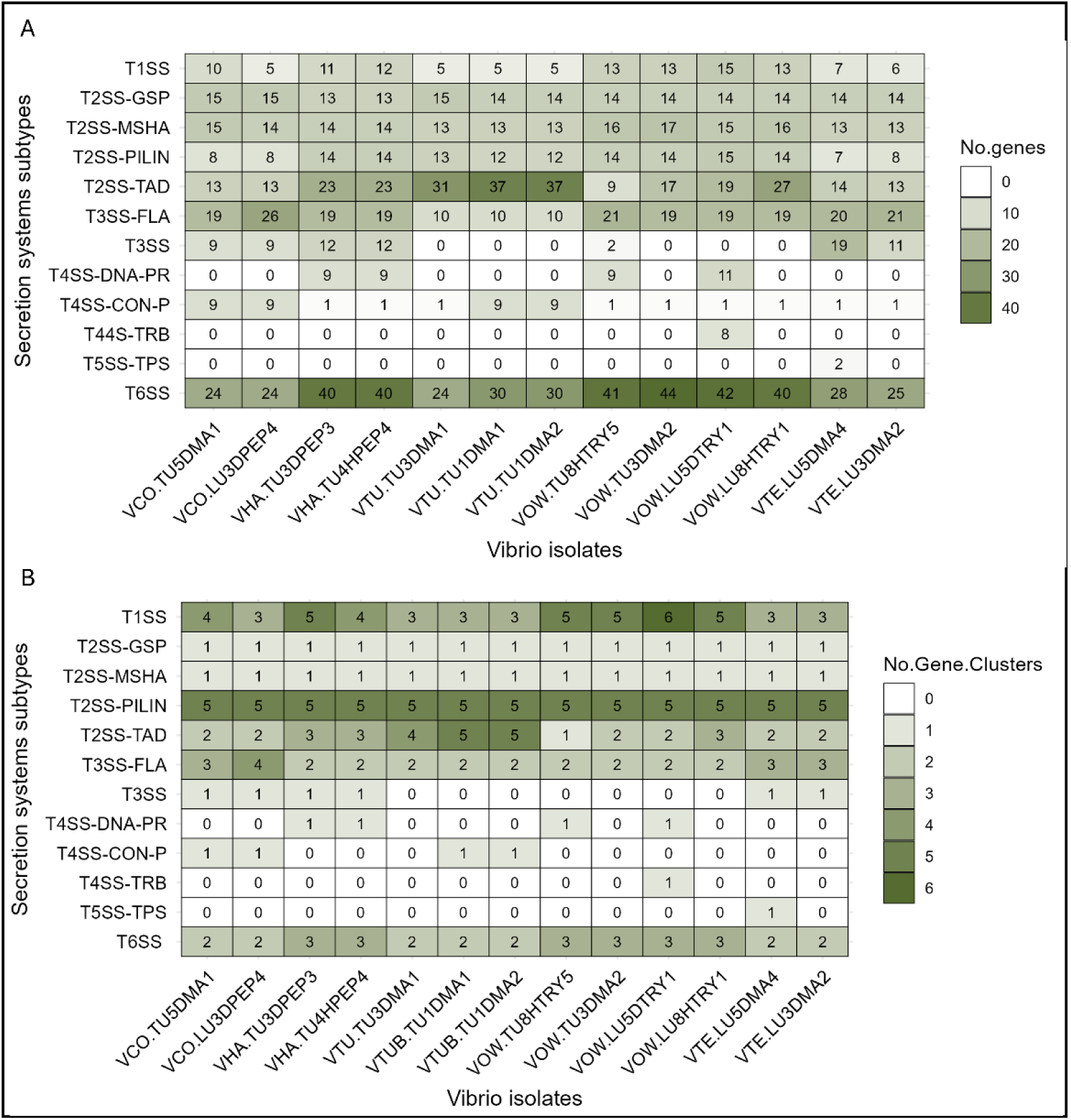
Distribution and organization of secretion system subtypes across thirteen *Vibrio* strains isolated from *Porites cylindrica*. (A) Total number of annotated genes assigned to each secretion system subtype based on KEGG BRITE classification (ko02044). (B) Number of discrete gene clusters identified per secretion system subtype within each genome. Secretion systems were grouped into twelve categories: type I secretion system (T1SS); type II secretion systems—general secretion pathway (T2SS-GSP), MSHA pilus biogenesis (T2SS-MSHA), pilin secretion and assembly proteins (T2SS-PILIN), and tight adherence export apparatus (T2SS-TAD); type III secretion systems—flagellar export apparatus (T3SS-FLA) and non-flagellar T3SS; type IV secretion systems—DNA and protein transfer (T4SS-DNA-PR), conjugative pilus assembly (T4SS-CON-P), and TRB system (T4SS-TRB); type V secretion system—two-partner secretion (T5SS-TPS); and type VI secretion system (T6SS). Strains are grouped by species: *Vibrio coralliilyticus* (VCO), *V. harveyi* (VHA), *V. tubiashii* (VTU), *V. owensii* (VOW), and *V. tetraodonis* (VTE).

**Figure 3.**
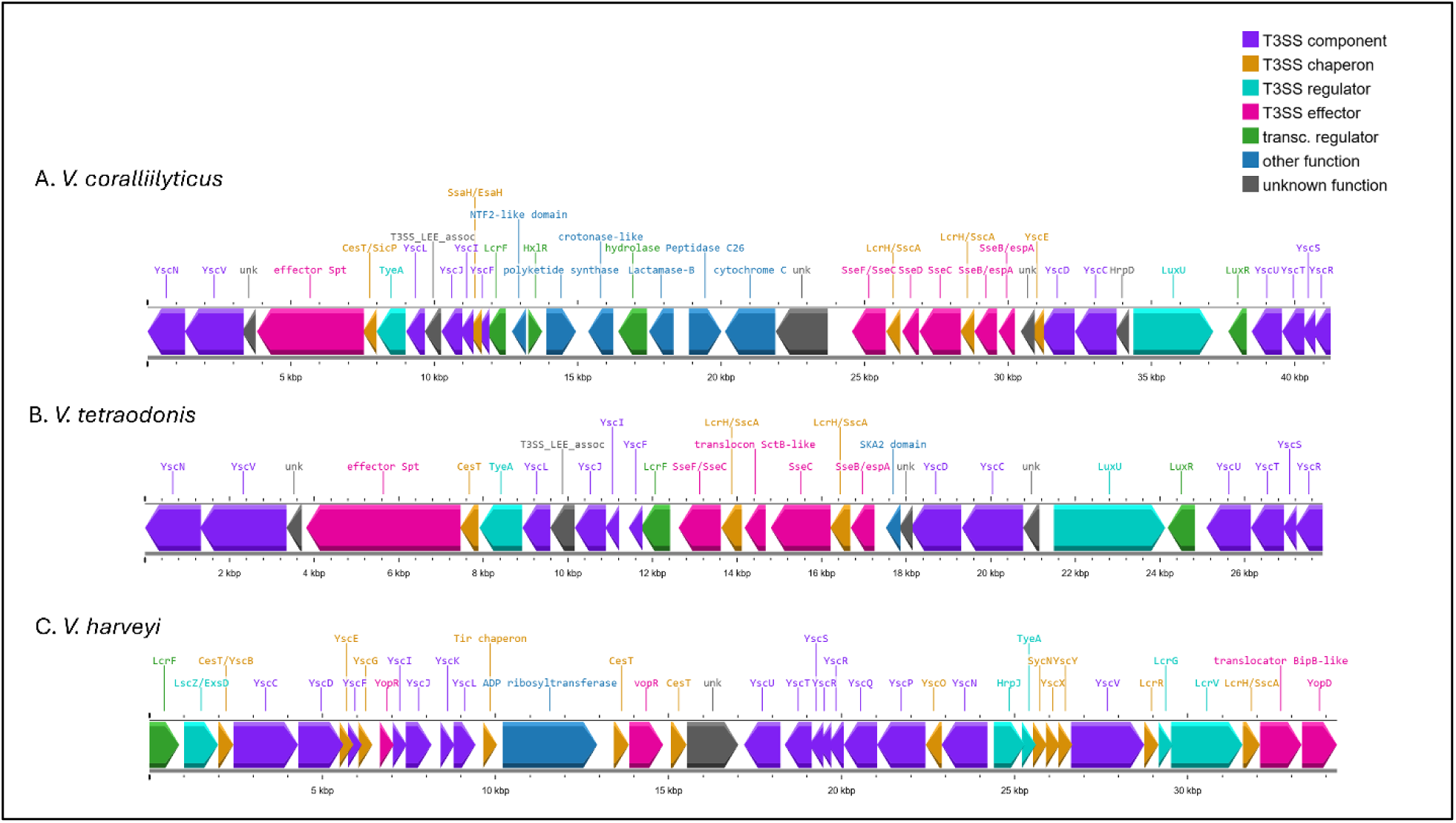
Comparative organization of type III secretion system (T3SS) gene clusters in three *Vibrio* species isolated from *Porites cylindrica*. (A) *Vibrio coralliilyticus*, (B) *V. tetraodonis*, and (C) *V. harveyi*. Arrows represent predicted open reading frames and indicate gene orientation. Colors denote functional categories: T3SS structural components (purple), chaperones (orange), regulators (green), predicted effectors (pink), transcriptional regulators (teal), other annotated functions (blue), and unknown/hypothetical proteins (gray). Gene names are shown above the arrows. Scale bars indicate genomic distance (kb). Synteny and gene-content differences among species highlight the diversification of T3SS architecture across coral-associated *Vibrio* lineages.

**Figure 4.**
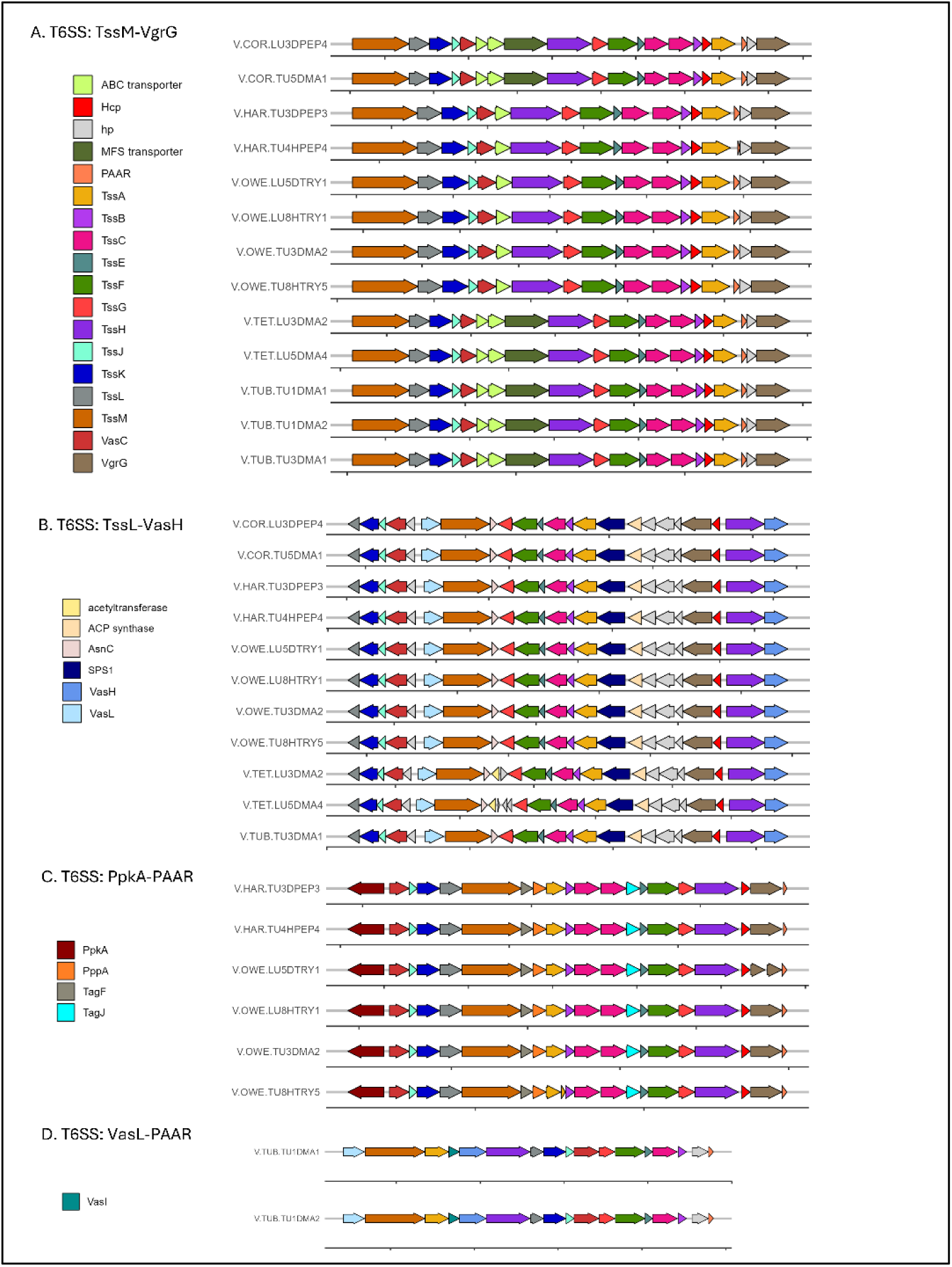
Comparative organization of Type VI secretion system (T6SS) subtypes across coral-associated *Vibrio* strains isolated from *Porites cylindrica*. Gene cluster architectures are shown for four structural T6SS subtypes: (A) TssM–VgrG, (B) TssL–VasH, (C) PpkA–PAAR, and (D) VasL–PAAR. Arrows represent predicted open reading frames and indicate gene orientation. Colors correspond to core structural components and accessory proteins as indicated in the legends for each panel. Scale bars represent genomic distance (kb). Strains are grouped by species: *Vibrio coralliilyticus* (V.COR), *V. harveyi* (V.HAR), *V. tubiashii* (V.TUB), *V. owensii* (V.OWE), and *V. tetraodonis* (V.TET). Conserved core structural genes are present across multiple species, whereas subtype-specific modules and accessory components vary among lineages

### Horizontal gene transfer elements in coral-associated *Vibrio* genomes

Horizontally acquired elements were identified across thirteen *Vibrio* genomes, including (i) plasmids (Table 3), (ii) prophages and episomal phages (Table 4), and (iii) genomic islands (GIs) and other mobile genetic elements (MGEs) predicted by IslandViewer4 and AlienHunter (Table 5). All detected plasmids and prophages were associated with strains isolated from diseased coral tissues, whereas genomic islands were present in both healthy- and disease-derived isolates.

**Table 3.**
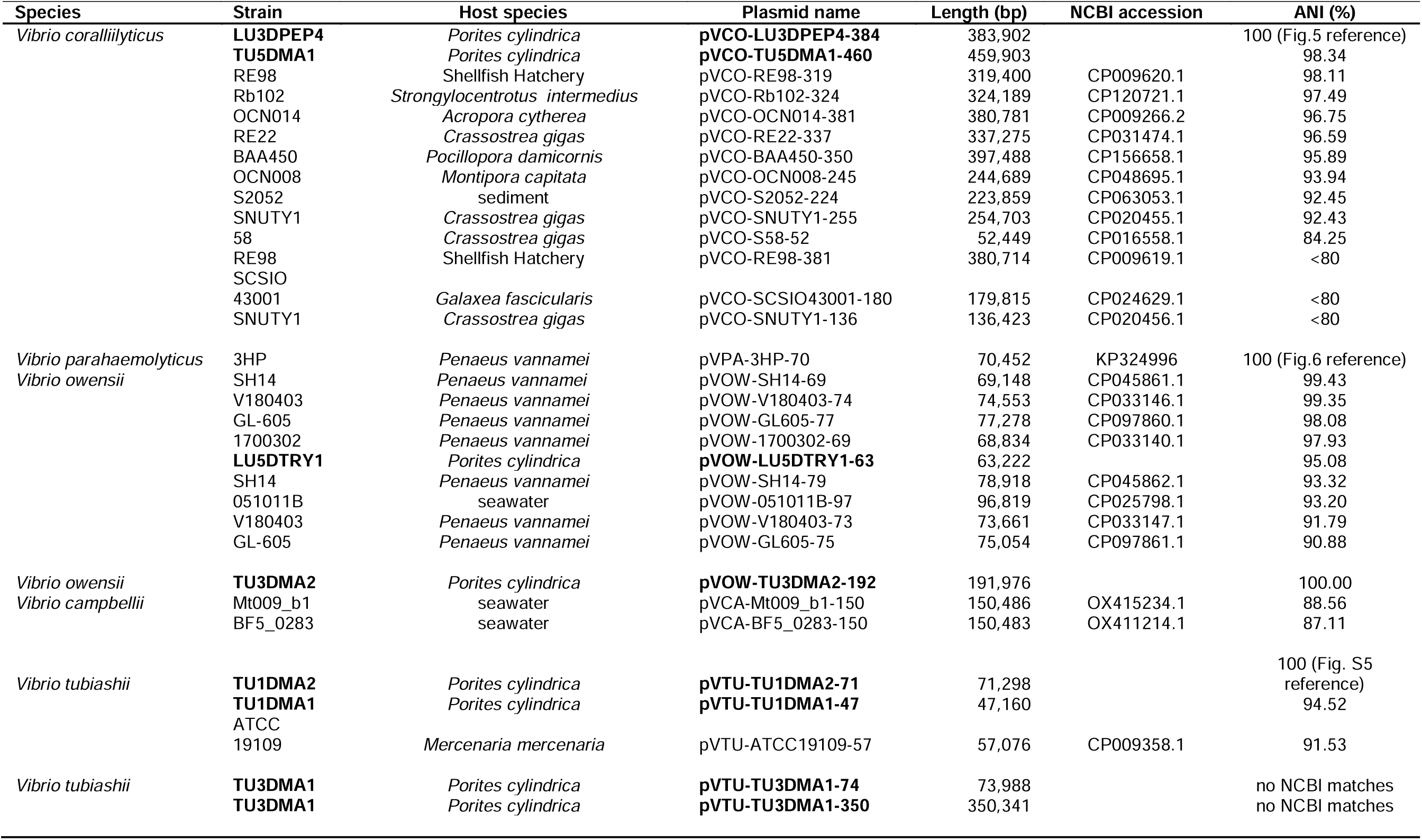
Comparative analysis of plasmids from *Vibrio* species associated with *Porites cylindrica* and other marine invertebrates. Plasmid size, NCBI accession numbers, and average nucleotide identity (ANI) relative to reference plasmids are shown. Plasmids sequenced in this study are indicated in bold. Plasmids are grouped according to ANI-based similarity to reference plasmids and correspond to comparative clusters visualized in (Figures 5, (6, and S5. Reference plasmids used for each comparison are indicated by 100% ANI values (see figure references in the ANI column).

**Table 4.** Genomic characteristics of intact prophages and episomal phages identified in Vibrio genomes using PHASTER. Phage location (ch: chromosome, p: plasmid), predicted genome length, PHASTER score (> 90 intact, 70–90 = questionable, < 70 = incomplete), and number of predicted proteins are shown.

| Species | Strain | Phage location | Phage name | Phage length | Phaster score | Number of proteins |
| --- | --- | --- | --- | --- | --- | --- |
| *Vibrio coralliilyticus* | LU3DPEP4 | p384:238548-2508780 | VCO.LU3DPEP4.p384.ph-12.3 | 12.3Kb | 136 | 20 |
| *Vibrio owensii* | TU3DMA2 | ch2:2175727-2296339 | VOW.TU3DMA2.ch2.ph1-120.6 | 120.6Kb | 140 | 121 |
| *Vibrio tetraodonis* | LU3DMA2 | ch1:2688905-2705315 | VAQ.LU3DMA2.ch1.ph1-16.4 | 16.4Kb | 110 | 24 |
|  | LU5DMA4 | ch1:4221842-4238366 | VAQ.LU5DMA4.ch1.ph1-16.5 | 16.5Kb | 110 | 23 |
|  | LU5DMA4 | ch1:3863947-3899017 | VAQ.LU5DMA4.ch1.ph2-35.0 | 35.0Kb | 120 | 49 |
| *Vibrio tubiashii* | TU1DMA2 | ch1:3443868-3471160 | VTU.TU1DMA2.ch1.ph1-27.2 | 27.2Kb | 100 | 35 |
|  | TU1DMA2 | ch1:1522352-1555632 | VTU.TU1DMA2.ch1.ph2-33.2 | 33.2Kb | 150 | 40 |
|  | TU3DMA1 | ch1:291652-328233 | VTU.TU3DMA1.ch1.ph1-36.5 | 36.5Kb | 110 | 51 |
|  | TU3DMA1 | ch1:600700-637305 | VTU.TU3DMA1.ch1.ph2-36.6 | 36.6Kb | 140 | 48 |
|  | TU3DMA1 | ch2:1263463-1303480 | VTU.TU3DMA1.ch2.ph3-40.0 | 40.0Kb | 130 | 48 |
|  | TU3DMA1 | p74:26-73851 | VTU.TU3DMA1.p74.ph4-73.8 | 73.8Kb | 150 | 105 |

**Table 5.** Distribution of virulence factors and secretion system genes within predicted and comparatively supported HGT regions. HGT-P: regions predicted as horizontally transferred using IslandViewer4 and AlienHunter. HGT-CS: predicted HGT regions with comparative support based on whole-chromosome BLAST alignments showing absence or discontinuity relative to closely related reference genomes (see Methods). Only HGT regions containing virulence factors or secretion system genes were evaluated for comparative support.

|  | *V. tubiashii* |  |  | *V. owensii* |  |  |  | *V. harveyi* |  | *V. coralliilyticus* |  | *V. tetraodonis* |  |
| --- | --- | --- | --- | --- | --- | --- | --- | --- | --- | --- | --- | --- | --- |
|  | TU3DMA1 | TU1DMA1 | TU1DMA2 | TU8HTRY5 | TU3DMA2 | LU5DTRY1 | LU8HTRY1 | TU3DPEP3 | TU4HPEP4 | TU5DMA1 | LU3DPEP4 | LU5DMA4 | LU3DMA2 |
| Virulence factors (total) | 8 | 8 | 8 | 7 | 11 | 8 | 8 | 6 | 6 | 14 | 17 | 11 | 12 |
| Virulence factors (HGT-P) | 1 | 2 | 4 | 1 | 3 | 2 | 2 | 1 | 1 | 5 | 1 | 7 | 4 |
| Virulence factors (HGT-CS) | 1 | 1 | 1 | 0 | 1 | 1 | 1 | 0 | 0 | 1 | 4 | 0 | 0 |
| T1SS (total) | 3 | 3 | 3 | 5 | 5 | 6 | 5 | 5 | 4 | 4 | 3 | 3 | 3 |
| T1SS (HGT-P) | 2 | 2 | 2 | 3 | 2 | 4 | 3 | 2 | 2 | 3 | 1 | 2 | 2 |
| T1SS (HGT-CS) | 1 | 1 | 1 | 1 | 0 | 1 | 1 | 1 | 0 | 1 | 0 | 0 | 2 |
| T2SS-TAD (total) | 4 | 5 | 5 | 1 | 2 | 2 | 3 | 3 | 3 | 2 | 2 | 2 | 2 |
| T2SS-TAD (HGT-P) | 2 | 4 | 4 | 1 | 1 | 2 | 3 | 2 | 2 | 2 | 1 | 1 | 1 |
| T2SS-TAD (HGT-CS) | 1 | 3 | 3 | 0 | 1 | 1 | 2 | 1 | 1 | 0 | 0 | 1 | 1 |
| T3SS (total) | 0 | 0 | 0 | 0 | 0 | 0 | 0 | 1 | 1 | 1 | 1 | 1 | 1 |
| T3SS (HGT-P) | 0 | 0 | 0 | 0 | 0 | 0 | 0 | 0 | 0 | 1 | 1 | 1 | 0 |
| T3SS (HGT-CP) | 0 | 0 | 0 | 0 | 0 | 0 | 0 | 0 | 0 | 1 | 1 | 0 | 0 |
| T4SS (total) | 0 | 0 | 0 | 1 | 0 | 1 | 0 | 1 | 1 | 0 | 0 | 0 | 0 |
| T4SS (HGT-P) | 0 | 0 | 0 | 1 | 0 | 1 | 0 | 1 | 1 | 0 | 0 | 0 | 0 |
| T4SS (HGT-CS) | 0 | 0 | 0 | 1 | 0 | 1 | 0 | 1 | 1 | 0 | 0 | 0 | 0 |
| T6SS (total) | 2 | 2 | 2 | 3 | 3 | 3 | 3 | 3 | 3 | 2 | 2 | 2 | 2 |
| T6SS (HGT-P) | 1 | 1 | 0 | 2 | 1 | 1 | 1 | 1 | 1 | 1 | 1 | 0 | 1 |
| T6SS (HGT-CS) | 1 | 1 | 1 | 0 | 0 | 0 | 0 | 1 | 1 | 0 | 0 | 0 | 0 |

#### Plasmid-mediated HGT

A total of 14 plasmids from 12 *V. coralliilyticus* strains originating from diverse hosts and geographic locations were compared. Ten plasmids exhibited >85% amino acid identity (Figure 5, Table 3). Plasmids varied in size and structure, including prophage-derived insertions. A phage-derived insertion encoding two zona occludens toxin (*zot*) genes was identified on plasmids from *V. coralliilyticus* strains LU3DPEP4 (this study) and BAA-450 (Figure 5). Across the set of ten plasmids, additional putative genes annotated as a cytotoxin MCF and a leucocidin domain-containing protein were detected (Figure 5). Conjugative transfer system genes were also present. BLAST comparisons indicated that the 63-kb plasmid from *V. owensii* strain LU5DTRY1 shared 95% amino acid identity with a ∼70-kb plasmid reported in pathogenic *V. parahaemolyticus* strains associated with acute hepatopancreatic necrosis disease (AHPND) in shrimp (Table 3). Comparative analyses showed that the LU5DTRY1 plasmid lacked *pirAB* toxin genes (Figure 6). No additional annotated toxin genes were detected, although the plasmid encoded a predicted secreted trypsin-like peptidase belonging to an unclassified subgroup within the S1 peptidase family (MEROPS). Two additional plasmids from *V. tubiashii* strains TU1DMA1 (47 kbp) and TU1DMA2 (71 kbp) shared >90% amino acid identity to the 57-kbp plasmid from *V. tubiashii* strain ATCC 19109 (Table 3; Figure S5). These plasmids did not contain any annotated virulence genes, though many encoded proteins were not functionally annotated (Figure S5).

**Figure 5.**
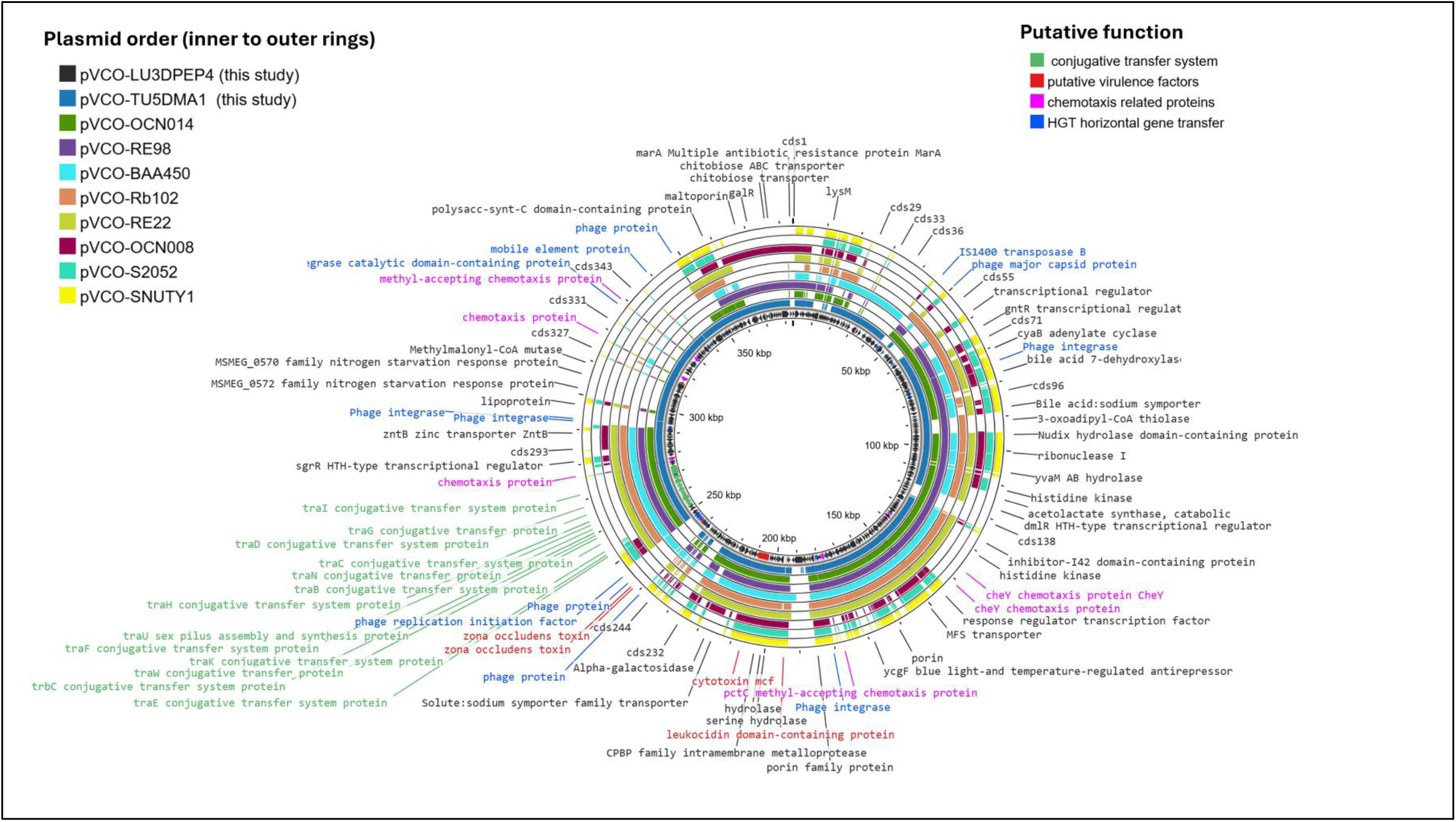
Comparative circular alignment of plasmids from ten *Vibrio coralliilyticus* strains, including two plasmids sequenced in this study (pVCO-LU3DPEP4 and pVCO-TU5DMA1) and eight reference plasmids from the NCBI database (pVCO-OCN014, pVCO-RE98, pVCO-BAA450, pVCO-Rb102, pVCO-RE22, pVCO-OCN008, pVCO-S2052, pVCO-SNUTY1; see Table 3 for strain information and accession numbers). Each concentric ring represents a single plasmid and is labeled with its corresponding plasmid name. Colored features indicate functional gene categories, including conjugative transfer systems (green), putative virulence factors (red), chemotaxis-related proteins (pink), and horizontal gene transfer–associated genes (blue).

**Figure 6.**
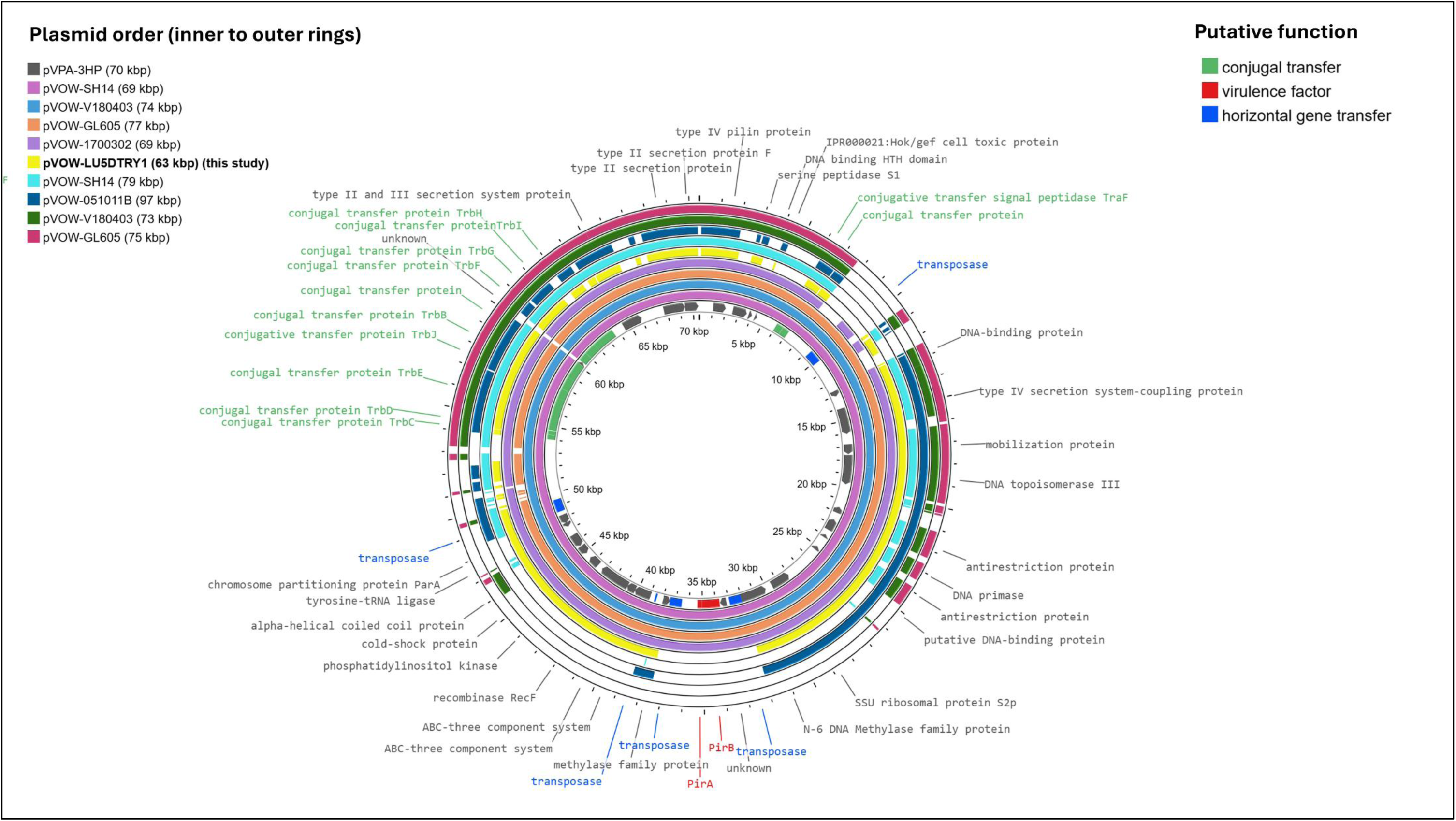
Comparative circular alignment of plasmids from 9 strains of *Vibrio owensii and* one strain of *V. parahaemolyticus*, including one plasmid from this study [pVOW-LU5DTRY1 (63 kbp)] and eight reference plasmids from the NCBI database [pVPA-3HP (70 kbp), pVOW-SH14 (69 kbp), pVOW-V180403 (74 kbp), pVOW-GL605 (77 kbp), pVOW-1700302 (69 kbp), pVOW-SH14 (79 kbp), pVOW-051011B (97 kbp), pVOW-V180403 (73 kbp), pVOW-GL605 (75 kbp); see Table 3 for strain information and accession numbers]. Each concentric ring represents a single plasmid and is labeled with its corresponding plasmid name. Colored features indicate functional gene categories, including conjugative transfer systems (green), putative virulence factors (red), and horizontal gene transfer–associated genes (blue).

#### Phage-mediated HGT

A total of 11 intact bacteriophages were identified across six *Vibrio* strains (Table 4; Figure 7; Supplementary Figure S4). Nine phages were integrated into host chromosomes, one was located within a plasmid (pVCO-LU3DPEP4-384, Table 3), and one existed as a free circular phage (V.TUB.ph-73.8, Table 4). This episomal phage was initially annotated as a plasmid (pVTU-TU3DMA1-74) but was subsequently recognized as a phage maintained episomally during lysogeny [58]. Prophage annotation identified virulence-associated genes in three phage regions. Accessory cholera toxin (*ace*) and zona occludens toxin (*zot*) genes were detected in prophages V.COR.ph-12.3 and V.OWE.ph-120.6 (Figure 7A, 7C). A prophage in *V. tetraodonis* (V.TET.ph-16.4) encoded a pectin lyase (Figure 7B).

**Figure 7.**
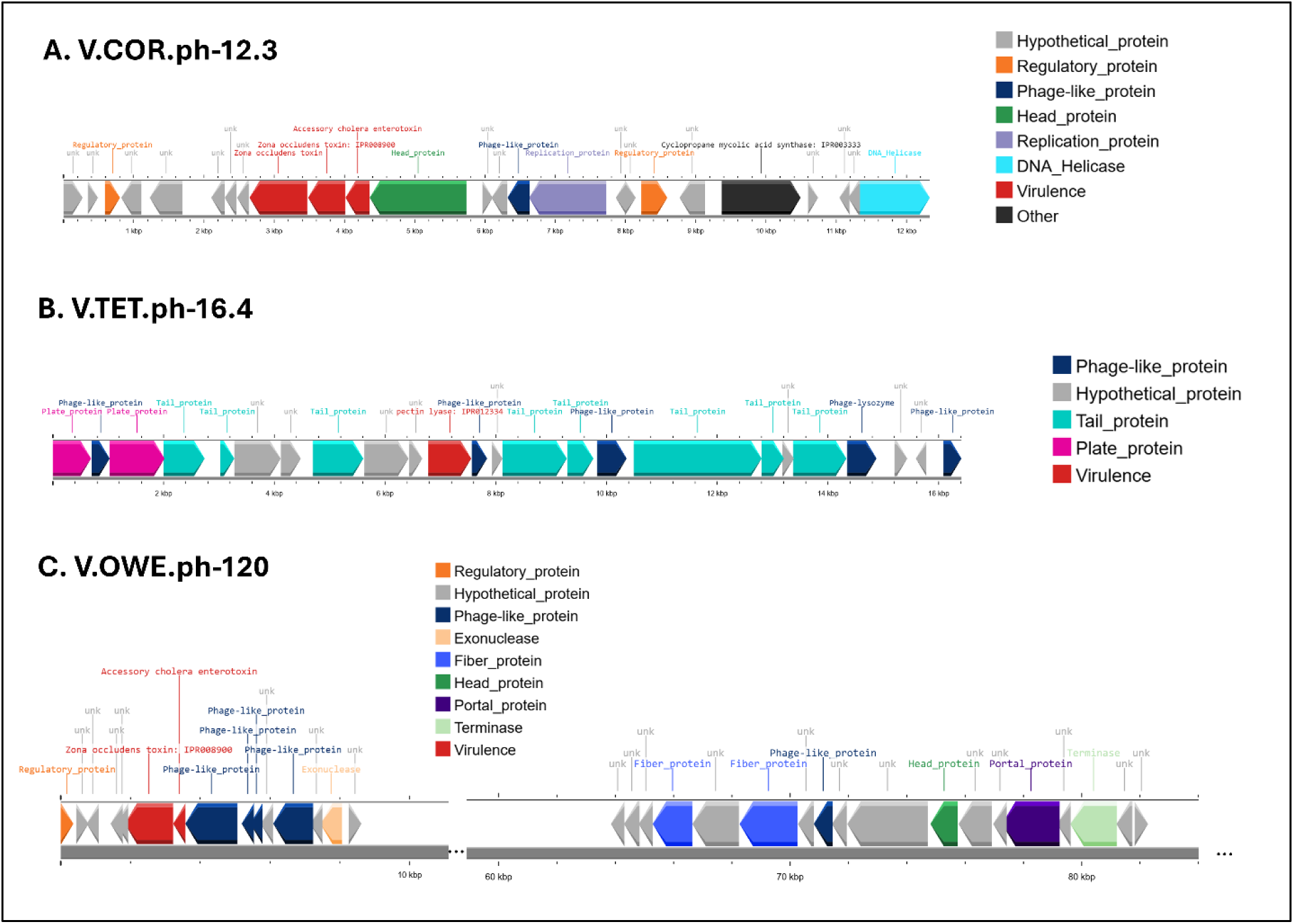
Virulence-associated prophage regions identified in coral-associated *Vibrio* genomes. Genomic organization of intact prophages carrying putative virulence-related genes in *Vibrio* strains isolated from *Porites cylindrica*. (A) V.COR.ph-12.3 from *V. coralliilyticus* containing accessory cholera enterotoxin (*ace*) and zona occludens toxin (*zot*). (B) V.TET.ph-16.4 from *V. tetraodonis* containing a pectin lyase gene. (C) V.OWE.ph-120.6 from *V. owensii* containing *ace* and *zot*. Arrows represent predicted open reading frames and indicate gene orientation. Colors denote functional categories as indicated in the legend (e.g., phage structural proteins, replication proteins, regulatory proteins, virulence-associated genes, hypothetical proteins). Scale bars indicate genomic distance (kb). Prophage genome locations within host chromosomes or plasmids are provided in Table 4.

#### Genomic islands and mobile genetic elements

Putative HGT regions were predicted using IslandViewer4 and AlienHunter (Table 5). IslandViewer4 identified between 2 and 89 GIs per chromosome, corresponding to 2–21% of chromosome length, whereas AlienHunter predicted 20–153 fragments per genome, covering 6–37% of each chromosome. Because both tools can generate false positives [59], only HGT regions containing virulence factors (Figure 1) and/or secretion system genes (Figure 2) were further evaluated using whole-chromosome alignments against reference genomes (Table 5; e.g., Figure 8). T4SS gene clusters identified in *V. harveyi* genomes were located within GIs (Figure 8). These islands contained at least two integrases and one relaxase, consistent with integrative and conjugative elements (ICEs) [60]. Although these ICEs lacked annotated toxin genes, they contained genes associated with adaptation (e.g., iron-scavenging and cold-shock proteins). In *V. owensii*, T4SS-bearing islands occurred at distinct genomic positions in TU8HTRY5 (large chromosome) and LU5DTRY1 (small chromosome), indicating variability in integration sites. T3SS gene clusters in the two *V. coralliilyticus* strains were located within regions homologous to a previously described pathogenicity island [35]. Comparative analysis indicated that all ten complete *V. coralliilyticus* genomes available in NCBI contained T3SS clusters (data not shown), whereas a closely related Vibrio sp. SCSIO 43001 (86% ANI to *V. coralliilyticus*) lacked the corresponding T3SS-containing island. Additional secretion system loci were also associated with predicted HGT regions (Table 5), including 15 of 36 T2SS-TAD clusters, 10 of 52 T1SS clusters, and several T6SS clusters. In *V. tubiashii,* strains differed in the T6SS subtype present within HGT regions (Table 5).

**Figure 8.**
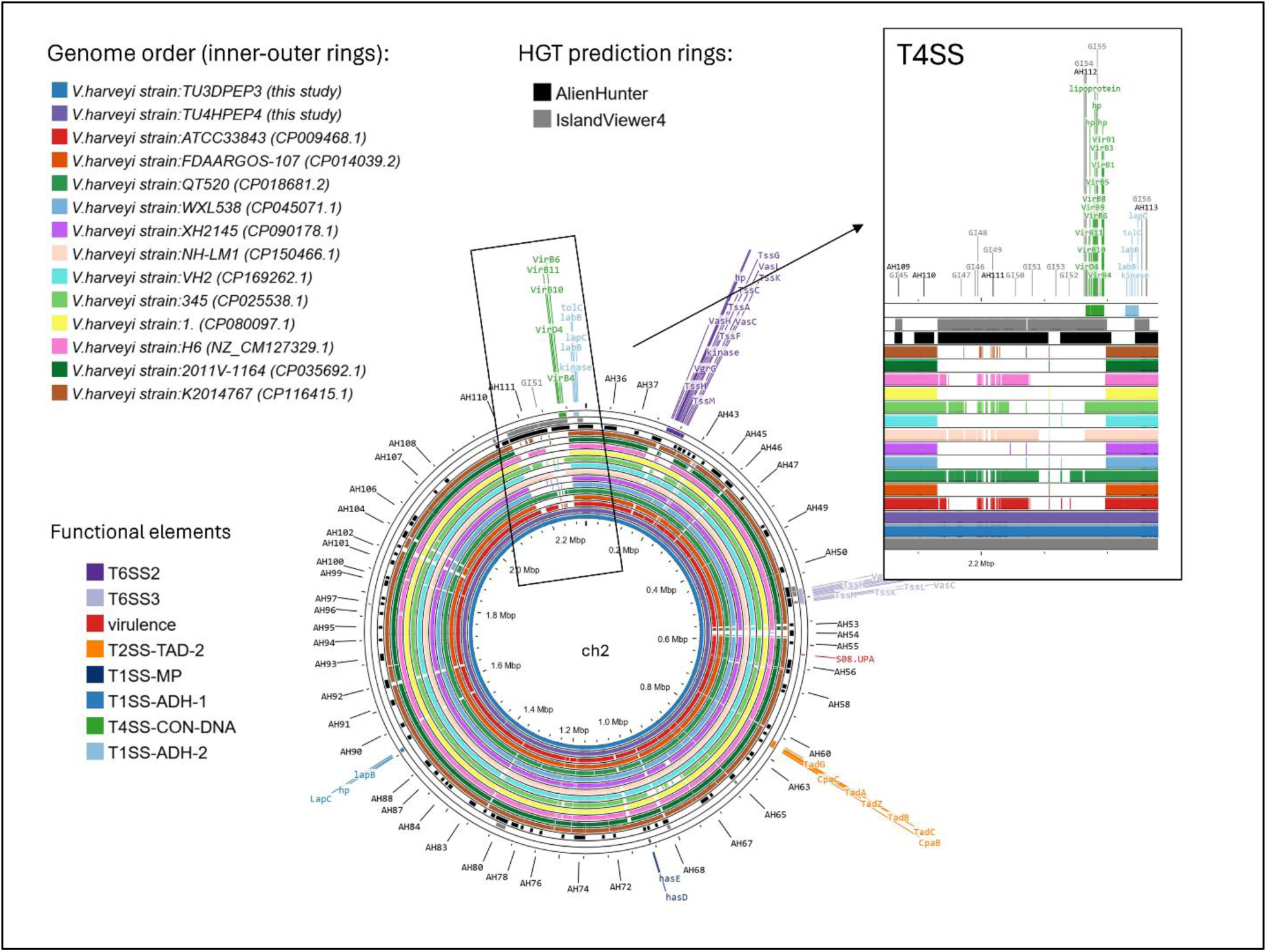
Comparative genomic context of a predicted horizontal gene transfer (HGT) region carrying a Type IV secretion system (T4SS) on chromosome 2 of *Vibrio harveyi*. Circular genome comparison generated using Proksee showing chromosome 2 (ch2) of *Vibrio harveyi* strains TU3DPEP3 and TU4HPEP4 (this study) as the innermost rings. The outer rings represent homologous regions from related *V. harveyi* reference genomes (NCBI accession number in parentheses). Colored segments indicate regions of sequence similarity identified by BLAST-based alignment. Predicted HGT regions are shown as separate rings: IslandViewer4 (gray) and AlienHunter (black). Overlapping predictions between the two tools indicate regions with stronger computational support for horizontal acquisition. Functional elements detected within the focal region are highlighted and color-coded, including T4SS components (green), T6SS clusters (purple), T2SS-TAD-2 (orange), T1SS subtypes (blue shades), and additional virulence-associated genes (red). Gene labels are shown for representative components within the enlarged inset panel. The insert displays a magnified view of the T4SS-containing genomic island, illustrating gene organization and correspondence with predicted HGT boundaries. Comparative absence or divergence of this region among reference genomes supports its potential horizontal acquisition.

## Discussion

### Distribution and genomic diversity of coral-associated *Vibrio*

In the current study, nearly half of the cultured bacterial isolates from *P. cylindrica* belonged to the genus *Vibrio*, underscoring the prominence of this group within the cultivable coral-associated microbiota. Among the five dominant *Vibrio* species isolated in this study, four—*V. owensii, V. tubiashii, V. coralliilyticus*, and *V. harveyi*—have previously been identified as coral pathogens [10, 14, 16, 19, 20]. In contrast, the group classified as *V. tetraodonis/aquimaris* has not been associated with coral disease or pathogenicity in marine animals to date [61, 62]. However, *Vibrio* isolates were not restricted to diseased tissue; they were also obtained from visually healthy colonies, but at much smaller numbers. Although culture-based isolation does not directly correspond to *in situ* abundance and may be influenced by growth conditions, the recovery of established coral pathogens from both healthy and diseased tissues is consistent with observations that several *Vibrio* species can be present in asymptomatic coral hosts and behave as opportunistic pathogens under changing environmental conditions [63–65]. In many coral diseases, a single obligate pathogen is not consistently identified, and outbreaks have been associated with shifts in host physiology and microbiome structure, as well as environmental stressors such as elevated temperature [66–68]. Within this framework, pathogenic potential may be broadly distributed among coral-associated *Vibrio* populations, with disease emergence reflecting stress-mediated activation or proliferation of virulence-capable taxa rather than the invasion of a discrete pathogenic lineage [64, 69]. The genomic architecture of the sequenced strains further highlights the diversity and plasticity of coral-associated *Vibrio*. Most genomes displayed the canonical two-chromosome structure typical of the genus, whereas one *V. tetraodonis* strain exhibited a single-chromosome configuration, potentially reflecting chromosomal fusion events similar to those described in *V. cholerae* [70, 71]. In addition, the presence of plasmids in multiple lineages indicates ongoing genetic exchange and the potential acquisition of accessory traits, as plasmid-mediated horizontal gene transfer has been shown to disseminate genes for adaptation, virulence, and other functions across bacterial populations [72, 73]. Collectively, these findings suggest that coral-associated *Vibrio* populations comprise a genetically heterogeneous assemblage characterized by conserved genomic architecture, structural variation, and mobile elements. This genomic diversity provides a foundation for exploring how virulence-associated genes, secretion systems, and horizontally acquired elements may contribute to opportunistic pathogenicity and coral disease dynamics.

### Conserved and lineage-specific virulence architectures in coral-associated *Vibrio*

Comparative analysis of virulence-associated genes across thirteen *Vibrio* strains representing five species and both healthy and disease-associated coral tissues revealed a layered virulence architecture, comprising broadly conserved extracellular enzymes and more restricted toxin and regulatory modules, consistent with the modular organization of virulence determinants described in many bacterial pathogens [74–76]. Lineage-restricted genes such as *vvhA, hlyA* homologs [33, 35, 77], cholera accessory toxins, and their associated regulators [31, 78, 79] as well as the *hcnABC* cluster [80–82] were confined to particular taxa, including *V. coralliilyticus* and *V. tetraodonis*. These genes represent potential lineage-specific enhancements of cytotoxicity or competitive ability, consistent with observations that horizontally acquired toxin modules and secretion systems can augment virulence phenotypes in bacterial pathogens [35, 74, 75]. Their restricted distribution suggests that while pathogenic potential may be broadly distributed within coral-associated *Vibrio* populations, certain lineages harbor additional modules that could intensify infection dynamics under favorable environmental or host conditions [64, 66, 69]. In contrast, extracellular proteases, including vibriolysins, collagenases, subtilisins [83–89], and the thermolabile hemolysin [90–92] were broadly distributed across taxa. Many of these enzymes have been implicated in virulence in other marine hosts. Yet, their universal or near-universal presence across strains from both healthy and diseased tissues suggests they likely serve dual ecological roles. In addition to contributing to tissue degradation during infection, these enzymes may facilitate nutrient acquisition from organic substrates within the coral-associated microbiome.

### Core and modular secretion architectures in coral-associated *Vibrio*

The comparative genomic analysis across the 13 strains reveals a layered secretion system architecture across coral-associated *Vibrio* genomes, characterized by a conserved motility and adhesion core alongside lineage-specific injectosome and competitive modules, consistent with the modular organization of bacterial secretion systems described across Gram-negative pathogens [93, 94]. Flagellar systems and conserved T2SS components were present in all strains, consistent with a shared ecological foundation for motility, surface attachment, and environmental persistence [24, 95, 96]. Variation in flagellar cluster organization and TAD copy number suggests differences in motility strategy and adhesion capacity among lineages [96, 97], but these systems were not restricted to isolates from diseased tissue. Greater lineage-level differentiation was observed in injectosome-type T3SS systems. T3SS3-like systems in *V. coralliilyticus* and *V. tetraodonis* and T3SS1-like systems in *V. harveyi* indicate diversification of host-targeting machinery [98–100]. However, T3SS was not exclusive to a single taxon, and predicted effectors were not confined to isolates from lesions, suggesting that host-targeting potential is modular rather than diagnostic of a specific pathogen [24, 93]. T4SS systems were frequently associated with plasmids, highlighting their role in horizontal gene transfer and genomic plasticity [101–103]. Their distribution across multiple species underscores the importance of mobile elements in shaping secretion system diversity and facilitating gene exchange [104]. T6SS exhibited both conserved and lineage-specific subtypes. The universal presence of TssM–VgrG suggests a shared competitive or host-interaction function [105, 106], while variation in additional subtypes indicates diversification of interbacterial and potentially host-targeting capacities [107]. Although *V. coralliilyticus* strains carried a broader repertoire of predicted anti-eukaryotic effectors [108], homologs were also present in other *Vibrio* isolates, reinforcing the distributed nature of potential virulence mechanisms. Overall, the diversity of secretion systems in coral-associated *Vibrio* reflects a modular, evolutionarily dynamic architecture [93, 94]. A conserved colonization toolkit is shared across taxa, while variable injectosome and competitive systems may modulate ecological interactions and pathogenic potential [24, 105]. Importantly, no single secretion system uniquely defined isolates recovered from diseased tissues, supporting an opportunistic model in which virulence capacity is distributed across multiple lineages and shaped by genomic plasticity rather than by a discrete obligate pathogen [64, 66, 69].

### HGT as a driver of virulence modularity and genomic plasticity in coral-associated *Vibrio*

Across these coral-associated *Vibrio* genomes, horizontally acquired elements—including plasmids, prophages/episomal phages, and genomic islands—represent major sources of accessory genetic variation, consistent with the recognized role of HGT in shaping *Vibrio* ecology and evolution [109–111]. The plasmid comparisons indicate that plasmids can be highly conserved within lineages (e.g., across multiple *V. coralliilyticus* strains) yet structurally dynamic due to insertions such as prophage-derived regions [58, 112]. Notably, only a subset of plasmids carried identifiable virulence-associated genes (e.g., *zot* insertions), suggesting that many plasmids may function primarily as vehicles for gene exchange and adaptation rather than direct determinants of pathogenicity [103, 113]. However, plasmid-mediated acquisition of virulence determinants is well documented in *Vibrio*, including toxin-bearing plasmids associated with AHPND in shrimp [38, 114–116], illustrating the potential for rapid phenotypic shifts following plasmid transfer [60]. Phage-associated loci contributed additional virulence-related traits, including *ace* and *zot* genes and a pectin lyase encoded within prophage regions. Prophages are well known to mediate the transfer of virulence genes and genomic innovation in *Vibrio* spp. and other pathogens [109, 111]. The identification of a prophage-associated pectin lyase is consistent with documented lateral acquisition of polysaccharide-degrading pathways in marine Gammaproteobacteria [117] and with the established virulence of pectic enzymes in plant-associated bacteria [118], suggesting potential ecological or host-associated functionality in marine systems. Beyond direct gene carriage, the presence of multiple intact prophages across strains highlights the potential for phage-mediated genome remodeling and gene movement within coral-associated microbial communities [113, 119, 120]. Notably, plasmids and prophages were detected exclusively in strains isolated from diseased coral tissues, whereas genomic islands were identified in both healthy- and disease-associated isolates. This pattern should be interpreted with caution, as the majority of sequenced genomes in this study originated from diseased samples, which may bias the observed distribution of mobile genetic elements. However, the consistent association of plasmids and phage-related elements with disease-derived strains, together with their enrichment in virulence-associated genes (e.g., zot, ace) and conjugative systems, is consistent with the established role of these elements in mediating pathogenicity and rapid adaptation in *Vibrio* [38, 109–111]. These observations suggest that mobile genetic elements may contribute to enhanced virulence potential in strains associated with diseased coral tissues. However, broader sampling will be required to determine whether the observed pattern reflects a consistent biological difference.

Genomic islands served as key integration platforms for secretion systems and other adaptive traits. The placement of T4SS clusters within ICE-like islands and the localization of T3SS clusters within pathogenicity-island homologs support the view that host interaction machinery in *Vibrio* can be gained, lost, or rearranged through horizontal transfer [28, 121, 122]. The variability in island location and content among closely related strains further indicates ongoing genomic flux, a hallmark of *Vibrio* eco-evolutionary dynamics [110]. While bioinformatic prediction of genomic islands can vary across detection tools [59], the repeated localization of secretion systems within predicted HGT regions strengthens the inference of horizontal acquisition. Taken together, these findings support a model in which *Vibrio* virulence potential is not encoded solely within the conserved core genome but is shaped by a modular accessory gene pool redistributed through HGT [109–111]. Such genomic plasticity provides an evolutionary mechanism for rapid shifts in host interaction strategies and may contribute to opportunistic disease dynamics in coral-associated *Vibrio* populations [24, 35].

## Conclusions

Comparative genomic analysis of five *Vibrio* species and thirteen strains isolated from healthy and WS–affected *P. cylindrica* revealed a modular virulence landscape distributed across multiple lineages. All strains possessed conserved extracellular proteases, including vibriolysins, collagenases, and subtilases, consistent with a shared ecological toolkit for protein degradation and host association. In contrast, cytolysins (e.g., *vvhA* and *hlyA* homologs) and phage-associated cholera-related toxins were restricted to particular taxa, reflecting lineage-specific acquisition of accessory virulence modules. Among the strains examined, *Vibrio coralliilyticus* exhibited the highest diversity of predicted virulence determinants, followed by select *V. tetraodonis* and *V. owensii* strains, whereas *V. harveyi* and *V. tubiashii* harbored fewer identifiable toxin modules. Rather than defining a single obligate pathogen, these patterns support a continuum of pathogenic potential in coral-associated *Vibrio*, in which conserved ecological functions are complemented by lineage-specific enhancements that may intensify infection under permissive environmental or host conditions. Substantial variation in secretion systems and mobile genetic elements further highlights the genomic plasticity of coral-associated *Vibrio*. Core colonization machinery, including flagellar systems and conserved T2SS components, was shared across taxa, whereas injectosome-type T3SS and conjugative T4SS systems exhibited lineage-level variability consistent with horizontal acquisition. The diversity of T6SS subtypes suggests both conserved competitive functions and diversification of interbacterial or host-targeting capacities. Prophages and plasmids contributed additional layers of genomic flexibility, reinforcing the role of horizontal gene transfer in redistributing virulence-associated modules and reshaping host-interaction potential. Plasmids and prophages were detected exclusively in strains isolated from diseased coral tissues in this dataset, whereas genomic islands were present in both healthy- and disease-associated isolates. While this pattern may reflect the greater number of genomes sequenced from diseased samples, it raises the possibility that these mobile elements contribute disproportionately to virulence potential in disease-associated *Vibrio*. Expanding genomic sampling across a larger number of isolates from both healthy and diseased corals will be essential to determine whether this pattern represents a robust biological signal. Together, these findings support a model in which virulence capacity in coral-associated *Vibrio* emerges from a flexible accessory gene pool superimposed upon a conserved ecological core. Pathogenic potential appears broadly distributed across lineages rather than confined to a single species, consistent with an opportunistic disease framework in which environmental stress, host condition, and genomic plasticity collectively influence infection dynamics. This study provides a comparative genomic foundation for understanding how modular virulence architectures and horizontal gene transfer shape coral–microbe interactions, and underscores the need for future work that integrates functional assays, transcriptomics, and longitudinal genomic monitoring to link gene content to virulence expression and disease emergence under changing environmental conditions.

## Data Availability

The sequencing data have been deposited in NCBI under the BioProject ID PRJNA1354586 and the BioSample accessions SAMN52948606-SAMN52948618.

## Authors’ contributions

Ewelina Rubin designed the research investigation, collected samples for bacterial culture, isolated the bacteria, performed molecular identification, prepared sequencing libraries, and assisted with genome sequencing. She analyzed all data, prepared the Figures and tables, and wrote the manuscript. Gaurav Shimpi led genome sequencing, assisted with data analysis, and reviewed and edited the manuscript. Héloïse Rouzé assisted with experimental design and sample collection and reviewed and edited the manuscript. Bastian Bentlage provided funding to complete the study and reviewed and edited the manuscript.

## Acknowledgements

We thank the University of Guam Marine Laboratory for logistical and laboratory support. The coral collection was made possible by permit number SCR-MPA-23-014, issued by the Division of Aquatic Wildlife Resources (DAWR) of the Department of Agriculture (DOAG).

## References

1. Visram S, Douglas AE. Resilience and acclimation to bleaching stressors in the scleractinian coral Porites cylindrica. J Exp Mar Bio Ecol 2007;349:35–44.

2. Mello Athayde MA. Present and future coral physiology of the resilient coral Porites cylindrica (Dana, 1846). https://espace.library.uq.edu.au/view/UQ:8f8e563 (2021).

3. Asoh K. Recovery of Porites cylindrica from the 2007-summer tissue loss at Shiraho, Ishigaki Island, Japan. *Galaxea*, Journal of Coral Reef Studies 2009;11:27–32.

4. Fitt WK, Gates RD, Hoegh-Guldberg O, Bythell JC, Jatkar A, et al. Response of two species of Indo-Pacific corals, Porites cylindrica and Stylophora pistillata, to short-term thermal stress: The host does matter in determining the tolerance of corals to bleaching. Journal of experimental marine biology and ecology 2009;373:102–110.

5. Burdick D, Brown V, Miller R. A report of the Comprehensive Long-term Coral Reef Monitoring at Permanent Sites on Guam project. https://repository.library.noaa.gov/view/noaa/29489 (2019, accessed May 13, 2024).

6. Burdick D, Raymundo L, Drake D, Hershberger A. A decade of change on Guam’s coral reefs. 170; University of Guam Marine Laboratory. https://www.uog.edu/_resources/files/ml/technical_reports/UOGML_TechRep170_GLTMP_2023.p df (2023).

7. Myers RL, Raymundo LJ. Coral disease in Micronesian reefs: a link between disease prevalence and host abundance. Dis Aquat Organ 2009;87:97–104.

8. Lozada-Misa P, Kerr A, Raymundo L. Contrasting Lesion Dynamics of White Syndrome among the scleractinian corals Porites spp. PLoS One 2015;10:e0129841.

9. Greene A, Donahue MJ, Caldwell JM, Heron SF, Geiger E, et al. Coral Disease Time Series Highlight Size-Dependent Risk and Other Drivers of White Syndrome in a Multi-Species Model. Frontiers in Marine Science;7. Epub ahead of print 2020. DOI: 10.3389/fmars.2020.601469.

10. Ben-Haim Y, Zicherman-Keren M, Rosenberg E. Temperature-regulated bleaching and lysis of the coral *Pocillopora damicornis* by the novel pathogen *Vibrio coralliilyticus*. Appl Environ Microbiol 2003;69:4236–4242.

11. Raymundo LJ, Andersen MD, Rouzé H. Coral restoration in a stressful environment: Disease, bleaching, and dysbiosis in Acropora aspera in Guam, Micronesia. iScience 2025;28:112244.

12. Bourne DG, Ainsworth TD, Willis BL. White syndromes of Indo-pacific corals. In: Diseases of Coral. Hoboken, NJ: John Wiley & Sons, Inc; 2015. pp. 300–315.

13. Morais J, Cardoso APLR, Santos BA. A global synthesis of the current knowledge on the taxonomic and geographic distribution of major coral diseases. Environmental Advances 2022;8:100231.

14. Séré MG, Tortosa P, Chabanet P, Quod J-P, Sweet MJ, et al. Identification of a bacterial pathogen associated with Porites white patch syndrome in the Western Indian Ocean. Mol Ecol 2015;24:4570–4581.

15. Burke S, Pottier P, Lagisz M, Macartney EL, Ainsworth T, et al. The impact of rising temperatures on the prevalence of coral diseases and its predictability: A global meta-analysis. Ecol Lett 2023;26:1466–1481.

16. Luna GM, Bongiorni L, Gili C, Biavasco F, Danovaro R. Vibrio harveyi as a causative agent of the White Syndrome in tropical stony corals. Environ Microbiol Rep 2010;2:120–127.

17. Sussman M, Willis BL, Victor S, Bourne DG. Coral pathogens identified for White Syndrome (WS) epizootics in the Indo-Pacific. PLoS One 2008;3:e2393.

18. Santos E de O, Alves N Jr, Dias GM, Mazotto AM, Vermelho A, et al. Genomic and proteomic analyses of the coral pathogen Vibrio coralliilyticus reveal a diverse virulence repertoire. ISME J 2011;5:1471–1483.

19. Ushijima B, Videau P, Burger AH, Shore-Maggio A, Runyon CM, et al. Vibrio coralliilyticus strain OCN008 is an etiological agent of acute Montipora white syndrome. Appl Environ Microbiol 2014;80:2102–2109.

20. Ushijima B, Smith A, Aeby GS, Callahan SM. Vibrio owensii induces the tissue loss disease Montipora white syndrome in the Hawaiian reef coral Montipora capitata. PLoS One 2012;7:e46717.

21. Zhenyu X, Shaowen K, Chaoqun H, Zhixiong Z, Shifeng W, et al. First characterization of bacterial pathogen, Vibrio alginolyticus, for Porites andrewsi White syndrome in the South China Sea. PLoS One 2013;8:e75425.

22. Rubin E, Raymundo L, Rouze H. White syndrome dynamics in Porites cylindrica: interactions among eutrophication, host structure, and microbial communities. bioRxiv. Epub ahead of print February 4, 2026. DOI: 10.64898/2026.02.04.703833.

23. Morris JG, Acheson D. Cholera and other types of vibriosis: a story of human pandemics and oysters on the half shell. Clin Infect Dis. https://academic.oup.com/cid/article-abstract/37/2/272/302936 (2003).

24. Baker-Austin C, Oliver JD, Alam M, Ali A, Waldor MK, et al. Vibrio spp. infections. Nat Rev Dis Primers 2018;4:8.

25. de Souza Valente C, Wan AHL. Vibrio and major commercially important vibriosis diseases in decapod crustaceans. J Invertebr Pathol 2021;181:107527.

26. Sampaio A, Silva V, Poeta P, Aonofriesei F. Vibrio spp.: Life strategies, ecology, and risks in a changing environment. Diversity (Basel). Epub ahead of print January 29, 2022. DOI: 10.3390/d14020097.

27. Aguirre-Guzman G, Mejia Ruiz H, Ascencio F. A review of extracellular virulence product of Vibrio species important in diseases of cultivated shrimp. Aquac Res 2004;35:1395–1404.

28. Kehlet-Delgado H, Häse CC, Mueller RS. Comparative genomic analysis of Vibrios yields insights into genes associated with virulence towards C. gigas larvae. BMC Genomics 2020;21:599.

29. Zhang X-H, He X, Austin B. Vibrio harveyi: a serious pathogen of fish and invertebrates in mariculture. Mar Life Sci Technol 2020;2:231–245.

30. Kim HJ, Jun JW, Giri SS, Chi C, Yun S, et al. Identification and Genome Analysis of Vibrio coralliilyticus Causing Mortality of Pacific Oyster (Crassostrea gigas) Larvae. Pathogens 2020;9:206.

31. Reidl J, Klose KE. *Vibrio cholerae*and cholera: out of the water and into the host. FEMS Microbiol Rev 2002;26:125–139.

32. Letchumanan V, Chan K-G, Lee L-H. Vibrio parahaemolyticus: a review on the pathogenesis, prevalence, and advance molecular identification techniques. Front Microbiol 2014;5:705.

33. Choi G, Choi SH. Complex regulatory networks of virulence factors in Vibrio vulnificus. Trends Microbiol 2022;30:1205–1216.

34. Manchanayake T, Salleh A, Amal MNA, Yasin ISM, Zamri-Saad M. Pathology and pathogenesis of Vibrio infection in fish: A review. Aquac Rep 2023;28:101459.

35. Kimes NE, Grim CJ, Johnson WR, Hasan NA, Tall BD, et al. Temperature regulation of virulence factors in the pathogen Vibrio coralliilyticus. ISME J 2012;6:835–846.

36. Weynberg KD, Voolstra CR, Neave MJ, Buerger P, van Oppen MJH. From cholera to corals: Viruses as drivers of virulence in a major coral bacterial pathogen. Sci Rep 2015;5:17889.

37. Lin H, Yu M, Wang X, Zhang X-H. Comparative genomic analysis reveals the evolution and environmental adaptation strategies of vibrios. BMC Genomics 2018;19:135.

38. Lee C-T, Chen I-T, Yang Y-T, Ko T-P, Huang Y-T, et al. The opportunistic marine pathogen *Vibrio parahaemolyticus* becomes virulent by acquiring a plasmid that expresses a deadly toxin. Proc Natl Acad Sci U S A 2015;112:10798–10803.

39. Galkiewicz JP, Kellogg CA. Cross-kingdom amplification using bacteria-specific primers: complications for studies of coral microbial ecology. Appl Environ Microbiol 2008;74:7828–7831.

40. Thompson FL, Gevers D, Thompson CC, Dawyndt P, Naser S, et al. Phylogeny and molecular identification of vibrios on the basis of multilocus sequence analysis. Appl Environ Microbiol 2005;71:5107–5115.

41. Quast C, Pruesse E, Yilmaz P, Gerken J, Schweer T, et al. The SILVA ribosomal RNA gene database project: improved data processing and web-based tools. Nucleic Acids Res 2013;41:D590–6.

42. Jain C, Rodriguez-R LM, Phillippy AM, Konstantinidis KT, Aluru S. High throughput ANI analysis of 90K prokaryotic genomes reveals clear species boundaries. Nat Commun 2018;9:5114.

43. Katoh K, Standley DM. MAFFT multiple sequence alignment software version 7: improvements in performance and usability. Mol Biol Evol 2013;30:772–780.

44. Trifinopoulos J, Nguyen L-T, von Haeseler A, Minh BQ. W-IQ-TREE: a fast online phylogenetic tool for maximum likelihood analysis. Nucleic Acids Res 2016;44:W232–5.

45. Nguyen L-T, Schmidt HA, von Haeseler A, Minh BQ. IQ-TREE: a fast and effective stochastic algorithm for estimating maximum-likelihood phylogenies. Mol Biol Evol 2015;32:268–274.

46. Kolmogorov M, Yuan J, Lin Y, Pevzner PA. Assembly of long, error-prone reads using repeat graphs. Nat Biotechnol 2019;37:540–546.

47. Simão FA, Waterhouse RM, Ioannidis P, Kriventseva EV, Zdobnov EM. BUSCO: assessing genome assembly and annotation completeness with single-copy orthologs. Bioinformatics 2015;31:3210–3212.

48. Aziz RK, Bartels D, Best AA, DeJongh M, Disz T, et al. The RAST Server: rapid annotations using subsystems technology. BMC Genomics 2008;9:75.

49. Liu B, Zheng D, Zhou S, Chen L, Yang J. VFDB 2022: a general classification scheme for bacterial virulence factors. Nucleic Acids Res 2022;50:D912–D917.

50. Kanehisa M, Sato Y, Morishima K. BlastKOALA and GhostKOALA: KEGG Tools for Functional Characterization of Genome and Metagenome Sequences. J Mol Biol 2016;428:726–731.

51. Rawlings ND, Barrett AJ, Bateman A. MEROPS: the peptidase database. Nucleic Acids Res 2010;38:D227–33.

52. Bertelli C, Laird MR, Williams KP, Simon Fraser University Research Computing Group, Lau BY, et al. IslandViewer 4: expanded prediction of genomic islands for larger-scale datasets. Nucleic Acids Res 2017;45:W30–W35.

53. Vernikos GS, Parkhill J. Interpolated variable order motifs for identification of horizontally acquired DNA: revisiting the Salmonella pathogenicity islands. Bioinformatics 2006;22:2196–2203.

54. Grant JR, Enns E, Marinier E, Mandal A, Herman EK, et al. Proksee: in-depth characterization and visualization of bacterial genomes. Nucleic Acids Res 2023;51:W484–W492.

55. Arndt D, Grant JR, Marcu A, Sajed T, Pon A, et al. PHASTER: a better, faster version of the PHAST phage search tool. Nucleic Acids Res 2016;44:W16–21.

56. Wilkins. . gggenes: Draw Gene Arrow Maps in “ggplot2”. R package version 0.5.0. https://wilkox.org/gggenes/ (2023, accessed 2025).

57. Zakaria D, Matsuda S, Iida T, Hayashi T, Arita M. Genome analysis identifies a novel type III secretion system (T3SS) category in Vibrio species. Microorganisms 2023;11:290.

58. Lan S-F, Huang C-H, Chang C-H, Liao W-C, Lin I-H, et al. Characterization of a new plasmid-like prophage in a pandemic Vibrio parahaemolyticus O3:K6 strain. Appl Environ Microbiol 2009;75:2659–2667.

59. da Silva Filho AC, Raittz RT, Guizelini D, De Pierri CR, Augusto DW, et al. Comparative analysis of genomic island prediction tools. Front Genet 2018;9:619.

60. Deng Y, Xu H, Su Y, Liu S, Xu L, et al. Horizontal gene transfer contributes to virulence and antibiotic resistance of Vibrio harveyi 345 based on complete genome sequence analysis. BMC Genomics 2019;20:761.

61. Franco A, Rückert C, Blom J, Busche T, Reichert J, et al. High diversity of Vibrio spp. associated with different ecological niches in a marine aquaria system and description of Vibrio aquimaris sp. nov. Syst Appl Microbiol 2020;43:126123.

62. Azevedo GPR, Mattsson HK, Lopes GR, Vidal L, Campeão M, et al. Vibrio tetraodonis sp. nov.: genomic insights on the secondary metabolites repertoire. Arch Microbiol 2021;203:399–404.

63. Rosenberg E, Koren O, Reshef L, Efrony R, Zilber-Rosenberg I. The role of microorganisms in coral health, disease and evolution. Nat Rev Microbiol 2007;5:355–362.

64. Bourne DG, Garren M, Work TM, Rosenberg E, Smith GW, et al. Microbial disease and the coral holobiont. Trends Microbiol 2009;17:554–562.

65. Sussman M, Mieog JC, Doyle J, Victor S, Willis BL, et al. Vibrio Zinc-Metalloprotease Causes Photoinactivation of Coral Endosymbionts and Coral Tissue Lesions. PLoS ONE 2009;4:e4511.

66. Harvell D, Jordán-Dahlgren E, Merkel S, Rosenberg E, Raymundo L, et al. Coral disease, environmental drivers, and the balance between coral and microbial associates. Oceanography 2007;20:172–195.

67. Vega Thurber R, Mydlarz LD, Brandt M, Harvell D, Weil E, et al. Deciphering Coral Disease Dynamics: Integrating Host, Microbiome, and the Changing Environment. Frontiers in Ecology and Evolution;8. Epub ahead of print 2020. DOI: 10.3389/fevo.2020.575927.

68. McDevitt-Irwin JM, Baum JK, Garren M, Vega Thurber RL. Responses of Coral-Associated Bacterial Communities to Local and Global Stressors. Frontiers in Marine Science;4. Epub ahead of print 2017. DOI: 10.3389/fmars.2017.00262.

69. Sweet MJ, Bulling MT. On the importance of the microbiome and pathobiome in coral health and disease. Front Mar Sci 2017;4:234438.

70. Chapman C, Henry M, Bishop-Lilly KA, Awosika J, Briska A, et al. Scanning the landscape of genome architecture of non-O1 and non-O139 Vibrio cholerae by whole genome mapping reveals extensive population genetic diversity. PLoS One 2015;10:e0120311.

71. Johnson SL, Khiani A, Bishop-Lilly KA, Chapman C, Patel M, et al. Complete genome assemblies for two single-chromosome Vibrio cholerae isolates, strains 1154-74 (serogroup O49) and 10432-62 (serogroup O27). Genome Announc;3. Epub ahead of print May 14, 2015. DOI: 10.1128/genomeA.00462-15.

72. Redondo-Salvo S, Fernández-López R, Ruiz R, Vielva L, de Toro M, et al. Pathways for horizontal gene transfer in bacteria revealed by a global map of their plasmids. Nat Commun 2020;11:3602.

73. Rodríguez-Beltrán J, DelaFuente J, León-Sampedro R, MacLean RC, San Millán Á. Beyond horizontal gene transfer: the role of plasmids in bacterial evolution. Nat Rev Microbiol 2021;19:347–359.

74. Hacker J, Kaper JB. Pathogenicity islands and the evolution of microbes. Annu Rev Microbiol 2000;54:641–679.

75. Dobrindt U, Hochhut B, Hentschel U, Hacker J. Genomic islands in pathogenic and environmental microorganisms. Nat Rev Microbiol 2004;2:414–424.

76. Rodriguez-Valera F, Martin-Cuadrado A-B, Rodriguez-Brito B, Pasić L, Thingstad TF, et al. Explaining microbial population genomics through phage predation. Nat Rev Microbiol 2009;7:828–836.

77. Debellis L, Diana A, Arcidiacono D, Fiorotto R, Portincasa P, et al. The Vibrio cholerae cytolysin promotes chloride secretion from intact human intestinal mucosa. PLoS One 2009;4:e5074.

78. Pérez-Reytor D, Jaña V, Pavez L, Navarrete P, García K. Accessory toxins of Vibrio pathogens and their role in epithelial disruption during infection. Front Microbiol 2018;9:2248.

79. Pruzzo C, Huq A, Colwell RR, Donelli G. Pathogenic Vibrio species in the marine and estuarine environment. In: Oceans and Health: Pathogens in the Marine Environment. Boston, MA: Springer US; 2006. pp. 217–252.

80. Askeland RA, Morrison SM. Cyanide production by Pseudomonas fluorescens and Pseudomonas aeruginosa. Appl Environ Microbiol 1983;45:1802–1807.

81. Laville J, Blumer C, Von Schroetter C, Gaia V, Défago G, et al. Characterization of the hcnABC gene cluster encoding hydrogen cyanide synthase and anaerobic regulation by ANR in the strictly aerobic biocontrol agent Pseudomonas fluorescens CHA0. J Bacteriol 1998;180:3187–3196.

82. Anderson RD, Roddam LF, Bettiol S, Sanderson K, Reid DW. Biosignificance of bacterial cyanogenesis in the CF lung. J Cyst Fibros 2010;9:158–164.

83. Shinoda S, Miyoshi S-I. Proteases produced by vibrios. Biocontrol Sci 2011;16:1–11.

84. Miyoshi S-I. Extracellular proteolytic enzymes produced by human pathogenic vibrio species. Front Microbiol 2013;4:339.

85. Vandeputte M, Kashem MA, Bossier P, Vanrompay D. *Vibrio* pathogens and their toxins in aquaculture: A comprehensive review. Rev Aquac 2024;16:1858–1878.

86. Galvis F, Barja J, Lemos M, Balado M. The vibriolysin-like protease VnpA and the collagenase ColA are required for full virulence of the bivalve mollusks pathogen Vibrio neptunius. Antibiotics (Basel);10. Epub ahead of print April 1, 2021. DOI: 10.3390/antibiotics10040391.

87. Ishihara M, Kawanishi A, Watanabe H, Tomochika K-I, Miyoshi S-I, et al. Purification of a serine protease of Vibrio parahaemolyticus and its characterization. Microbiol Immunol 2002;46:298–303.

88. Hasegawa H, Lind EJ, Boin MA, Häse CC. The extracellular metalloprotease of Vibrio tubiashii is a major virulence factor for pacific oyster (Crassostrea gigas) larvae. Appl Environ Microbiol 2008;74:4101–4110.

89. Lee C-Y, Cheng M-F, Yu M-S, Pan M-J. Purification and characterization of a putative virulence factor, serine protease, from Vibrio parahaemolyticus. FEMS Microbiol Lett 2002;209:31–37.

90. Wang S-X, Zhang X-H, Zhong Y-B, Sun B-G, Chen J-X. Genes encoding the Vibrio harveyi haemolysin (VHH)/thermolabile haemolysin (TLH) are widespread in vibrios. Wei Sheng Wu Xue Bao 2007;47:874–881.

91. Jia A, Woo NYS, Zhang X-H. Expression, purification, and characterization of thermolabile hemolysin (TLH) from Vibrio alginolyticus. Dis Aquat Organ 2010;90:121–127.

92. Taniguchi H, Kubomura S, Hirano H, Mizue K, Ogawa M, et al. Cloning and characterization of a gene encoding a new thermostable hemolysin from Vibrio parahaemolyticus. FEMS Microbiol Lett 1990;55:339–345.

93. Green ER, Mecsas J. Bacterial Secretion Systems: An Overview. Microbiol Spectr;4. Epub ahead of print February 2016. DOI: 10.1128/microbiolspec.VMBF-0012-2015.

94. Costa TRD, Felisberto-Rodrigues C, Meir A, Prevost MS, Redzej A, et al. Secretion systems in Gram-negative bacteria: structural and mechanistic insights. Nat Rev Microbiol 2015;13:343–359.

95. Echazarreta MA, Klose KE. Vibrio flagellar synthesis. Front Cell Infect Microbiol 2019;9:131.

96. McCarter LL. Dual flagellar systems enable motility under different circumstances. J Mol Microbiol Biotechnol 2004;7:18–29.

97. Meron D, Efrony R, Johnson WR, Schaefer AL, Morris PJ, et al. Role of flagella in virulence of the coral pathogen Vibrio coralliilyticus. Appl Environ Microbiol 2009;75:5704–5707.

98. Park K-S, Ono T, Rokuda M, Jang M-H, Okada K, et al. Functional characterization of two type III secretion systems of Vibrio parahaemolyticus. Infect Immun 2004;72:6659–6665.

99. Miller KA, Tomberlin KF, Dziejman M. Vibrio variations on a type three theme. Curr Opin Microbiol 2019;47:66–73.

100. Hajra D, Nair AV, Chakravortty D. An elegant nano-injection machinery for sabotaging the host: Role of Type III secretion system in virulence of different human and animal pathogenic bacteria. Phys Life Rev 2021;38:25–54.

101. Christie PJ. The mosaic type IV secretion systems. EcoSal Plus;7. Epub ahead of print October 2016. DOI: 10.1128/ecosalplus.ESP-0020-2015.

102. Backert S, Meyer TF. Type IV secretion systems and their effectors in bacterial pathogenesis. Curr Opin Microbiol 2006;9:207–217.

103. Adamczyk M, Jagura-Burdzy G. Spread and survival of promiscuous IncP-1 plasmids. Acta Biochim Pol 2003;50:425–453.

104. Rahmani A, Delavat F, Lambert C, Le Goic N, Dabas E, et al. Implication of the type IV secretion system in the pathogenicity of Vibrio tapetis, the etiological agent of Brown Ring Disease affecting the Manila clam Ruditapes philippinarum. Front Cell Infect Microbiol 2021;11:634427.

105. Russell AB, Peterson SB, Mougous JD. Type VI secretion system effectors: poisons with a purpose. Nat Rev Microbiol 2014;12:137–148.

106. Crisan CV, Hammer BK. The Vibrio cholerae type VI secretion system: toxins, regulators and consequences. Environ Microbiol 2020;22:4112–4122.

107. Jana B, Keppel K, Fridman CM, Bosis E, Salomon D. Multiple T6SSs, mobile auxiliary modules, and effectors revealed in a systematic analysis of the Vibrio parahaemolyticus pan-genome. mSystems 2022;7:e0072322.

108. Mass S, Cohen H, Gerlic M, Ushijima B, van Kessel JC, et al. A T6SS in the coral pathogen*Vibrio coralliilyticus*secretes an arsenal of anti-eukaryotic effectors and contributes to virulence. bioRxiv 2024;2024.03.20.584600.

109. Hazen TH, Pan L, Gu J-D, Sobecky PA. The contribution of mobile genetic elements to the evolution and ecology of Vibrios: Vibrio HGT. FEMS Microbiol Ecol 2010;74:485–499.

110. Le Roux F, Blokesch M. Eco-evolutionary dynamics linked to horizontal gene transfer in vibrios. Annu Rev Microbiol 2018;72:89–110.

111. Brito IL. Examining horizontal gene transfer in microbial communities. Nat Rev Microbiol 2021;19:442–453.

112. Hazen TH, Wu D, Eisen JA, Sobecky PA. Sequence characterization and comparative analysis of three plasmids isolated from environmental Vibrio spp. Appl Environ Microbiol 2007;73:7703–7710.

113. Carr VR, Shkoporov A, Hill C, Mullany P, Moyes DL. Probing the mobilome: Discoveries in the dynamic microbiome. Trends Microbiol 2021;29:158–170.

114. Dong X, Chen J, Song J, Wang H, Wang W, et al. Evidence of the horizontal transfer of pVA1-type plasmid from AHPND-causing V. campbellii to non-AHPND V. owensii. Aquaculture 2019;503:396–402.

115. Dong X, Song J, Chen J, Bi D, Wang W, et al. Conjugative transfer of the pVA1-type Plasmid carrying the pirAB vp genes results in the formation of new AHPND-causing Vibrio. Front Cell Infect Microbiol 2019;9:195.

116. Xiao J, Liu L, Ke Y, Li X, Liu Y, et al. Shrimp AHPND-causing plasmids encoding the PirAB toxins as mediated by pirAB-Tn903 are prevalent in various Vibrio species. Sci Rep 2017;7:42177.

117. Hehemann J-H, Van Truong L, Unfried F, Welsch N, Kabisch J, et al. Aquatic adaptation of a laterally acquired pectin degradation pathway in marine gammaproteobacteria. Environ Microbiol 2017;19:2320–2333.

118. Herron SR, Benen JA, Scavetta RD, Visser J, Jurnak F. Structure and function of pectic enzymes: virulence factors of plant pathogens. Proc Natl Acad Sci U S A 2000;97:8762–8769.

119. Dahlman S, Avellaneda-Franco L, Barr JJ. Phages to shape the gut microbiota? Curr Opin Biotechnol 2021;68:89–95.

120. Lin DM, Lin HC. A theoretical model of temperate phages as mediators of gut microbiome dysbiosis. F1000Res 2019;8:997.

121. Morita M, Yamamoto S, Hiyoshi H, Kodama T, Okura M, et al. Horizontal gene transfer of a genetic island encoding a type III secretion system distributed in Vibrio cholerae. Microbiol Immunol 2013;57:334–339.

122. Jerez SA, Plaza N, Bravo V, Urrutia IM, Blondel CJ. Vibrio type III secretion system 2 is not restricted to the Vibrionaceae and encodes differentially distributed repertoires of effector proteins. Microb Genom;9. Epub ahead of print April 2023. DOI: 10.1099/mgen.0.000973.

