## Supplementary for "Comparative genomics reveals modular virulence repertoires and extensive horizontal gene transfer in *Vibrio* species associated with white syndrome of *Porites cylindrica*"

**Supplementary Table S1**: Number and taxonomic classification of bacterial isolates obtained from healthy and diseased tissue of *Porites cylindrica* samples from two reef locations

| | | Luminao | | Tumon Bay | |
|---|---|---|---|---|---|
| | | Healthy | Diseased | Healthy | Diseased |
| Gammaproteobacteria | Vibrionaceae | 4 | 21 | 8 | 25 |
| Alphaproteobacteria | Rhodobacteraceae | 15 | 15 | 0 | 0 |
| Alphaproteobacteria | Kiloniellaceae | 0 | 9 | 0 | 0 |
| Alphaproteobacteria | Eilatimonas | 2 | 0 | 0 | 0 |
| Alphaproteobacteria | Rhizobiaceae | 1 | 0 | 0 | 0 |
| Alphaproteobacteria | Stappiaceae | 2 | 0 | 5 | 0 |
| Bacteroidia | Cyclobacteriaceae | 0 | 1 | 0 | 0 |
| Bacteroidia | Flavobacteriaceae | 3 | 0 | 2 | 0 |
| Actinobacteria | Micrococcales | 0 | 2 | 0 | 0 |

**Supplementary Table S2** Average nucleotide identity (ANI) values between the 13 genomes sequenced in this study and the type strain genomes of their closest phylogenetic relatives identified by maximum-likelihood analysis.

| Reference species | Reference NCBI assembly accession | Isolate name -this study | ANI (%) | Reference genome fragments | Query genome fragments |
|---|---|---|---|---|---|
| *Vibrio owensii* | GCF_002021755.1 | LU5DTRY1 | 96.9907 | 1963 | 1846 |
| *Vibrio owensii* | GCF_002021755.1 | TU3DMA2 | 97.0455 | 1963 | 1848 |
| *Vibrio owensii* | GCF_002021755.1 | LU8HTRY1 | 97.0439 | 1963 | 1855 |
| *Vibrio owensii* | GCF_002021755.1 | TU8HTRY5 | 97.0232 | 1963 | 1843 |
| *Vibrio harveyi* | GCF_030060435.1 | TU3DPEP3 | 98.5397 | 1956 | 1793 |
| *Vibrio harveyi* | GCF_030060435.1 | TU4HPEP4 | 98.4822 | 1956 | 1801 |
| *Vibrio coralliilyticus* | GCF_029541605.1 | LU3DPEP4 | 97.2075 | 1918 | 1768 |
| *Vibrio coralliilyticus* | GCF_029541605.1 | TU5DMA1 | 97.2853 | 1918 | 1777 |
| *Vibrio tubiashii* | GCF_000772105.1 | TU1DMA1 | 95.6602 | 1843 | 1500 |
| *Vibrio tubiashii* | GCF_000772105.1 | TU3DMA1 | 95.7478 | 1843 | 1493 |
| *Vibrio tubiashii* | GCF_000772105.1 | TU1DMA2 | 95.6083 | 1843 | 1498 |
| *Vibrio aquimaris* | GCF_009363415.1 | LU3DMA2 | 94.6136 | 1499 | 1291 |
| *Vibrio aquimaris* | GCF_009363415.1 | LU5DMA4 | 94.8863 | 1499 | 1345 |
| *Vibrio tetraodonis* | GCF_003350295.1 | LU3DMA2 | 95.1874 | 1435 | 1279 |
| *Vibrio tetraodonis* | GCF_003350295.1 | LU5DMA4 | 95.4677 | 1435 | 1299 |

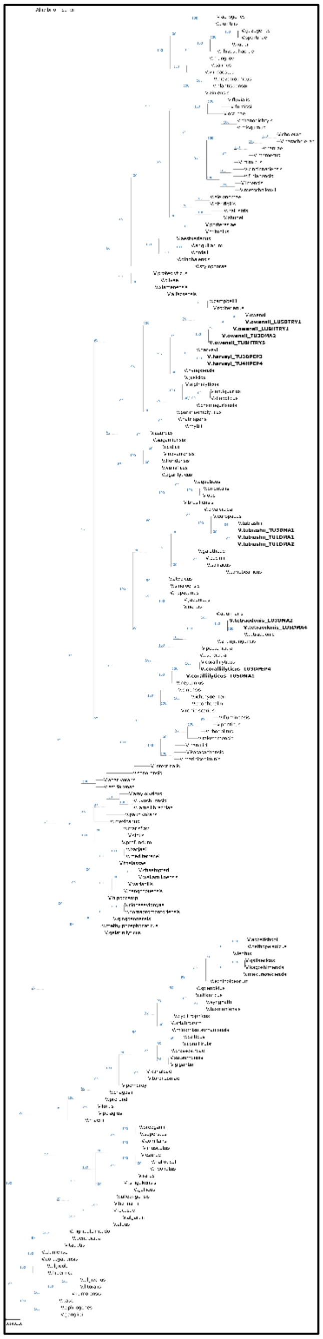

**Supplementary Figure S1.** Maximum-likelihood phylogeny based on a concatenated alignment of eight housekeeping genes (*ftsZ, gap, gyrB, pyrH, recA, topA, rpoA,* and *mreB*) from 189 taxa, including 175 publicly available Vibrio reference genomes, 13 isolates generated in this study, and one outgroup taxon. Gene alignments were concatenated into a single supermatrix and analyzed using IQ-TREE. Node support values are shown at each node based on ultrafast bootstrap support (%) (UFBoot). The tree is rooted using the designated outgroup. Scale bar indicates substitutions per site.

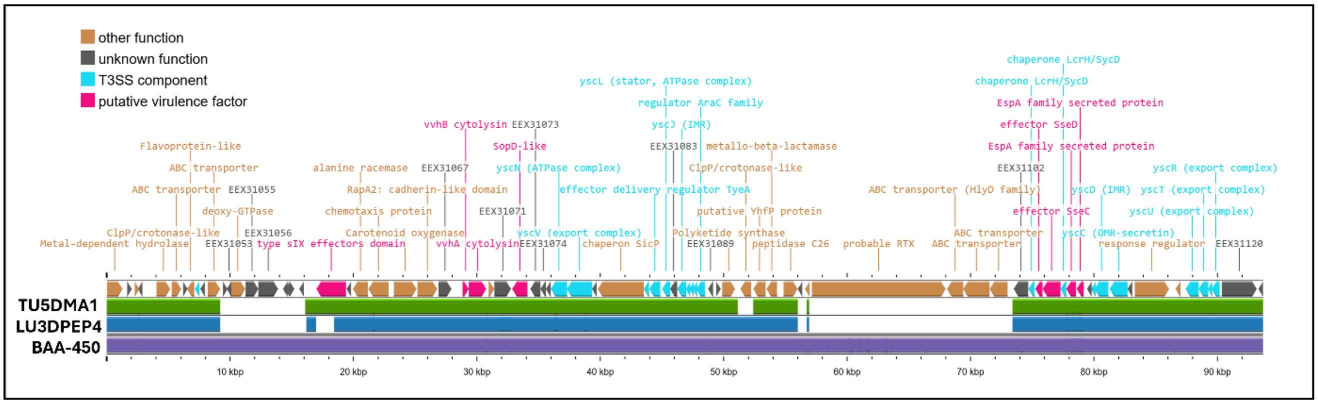


**Supplementary Figure S2. Comparative BLAST alignment of *Vibrio coralliilyticus* strains to the CP-1 pathogenicity island.** BLAST-based genomic comparison of *Vibrio coralliilyticus* strains LU3DPEP4 and TU5DMA1 (this study) against the CP-1 pathogenicity island from *V. coralliilyticus* strain BAA-450 (Kimes et al., 2012). The top panel shows gene organization of the CP-1 island (reference), with genes color-coded by functional category: T3SS components (cyan), putative virulence factors (magenta), other annotated functions (brown), and hypothetical proteins (gray). Horizontal alignment tracks below indicate regions of nucleotide similarity between the reference CP-1 island (BAA-450) and the genomes of LU3DPEP4 and TU5DMA1. Colored bars represent BLAST matches meeting the applied similarity thresholds, illustrating conservation and structural variation across strains. Conserved T3SS structural genes (e.g., yscC, yscN, yscT, yscU) and associated effector and regulatory elements are present in both strains, although differences in gene content and synteny are evident, suggesting partial conservation of the CP-1 island among *V. coralliilyticus* isolates.

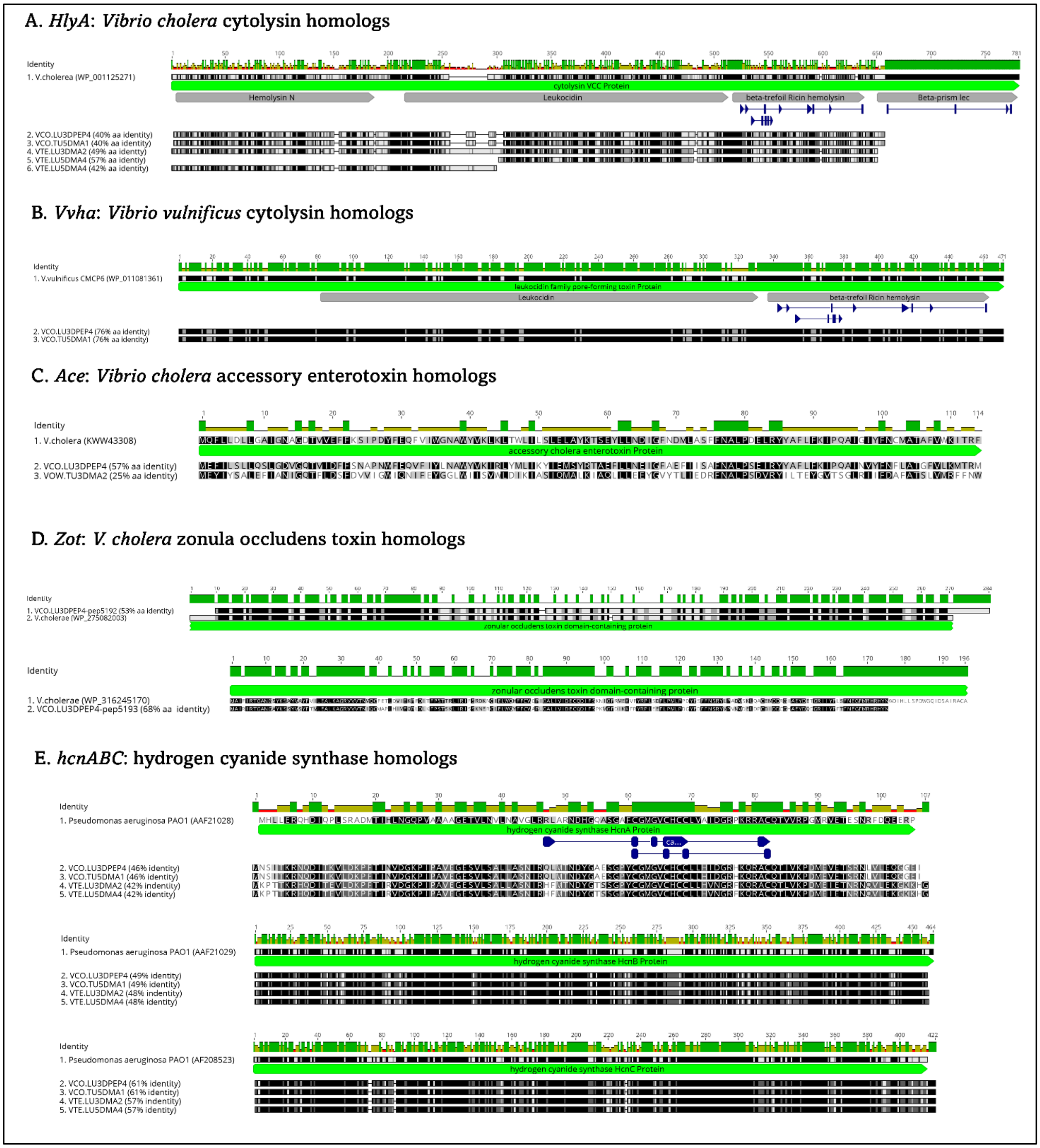


**Supplementary Figure S3. Amino acid alignments of selected virulence-associated proteins identified in *Vibrio* isolates from this study.** Multiple sequence alignments comparing representative virulence-associated proteins identified in this study with reference toxins from established pathogens. Percent amino acid identity relative to reference sequences is indicated for each homolog. (A) HlyA homologs aligned to *Vibrio cholerae* hemolysin (WP_001125271). (B) *VvhA* homologs aligned to *Vibrio vulnificus* cytolysin (WP_011081361). (C) Ace (accessory cholera enterotoxin) homologs aligned to *V. cholerae* Ace (KWW43308). (D) *Zot*, zonula occludens toxin homologs aligned to *V. cholerae* (WP_275082003, WP_316245170). (E) *HcnABC* hydrogen cyanide synthase subunits aligned to reference sequences from *Pseudomonas aeruginosa* PAO1 (AAF21028, AAF21029A, AF21030). Conserved domains annotated in reference proteins are shown above each alignment (e.g., leukocidin family pore-forming toxin domain, beta-trefoil ricin-like hemolysin domain). Green bars indicate conserved regions, and black/gray shading represents amino acid conservation across aligned sequences.

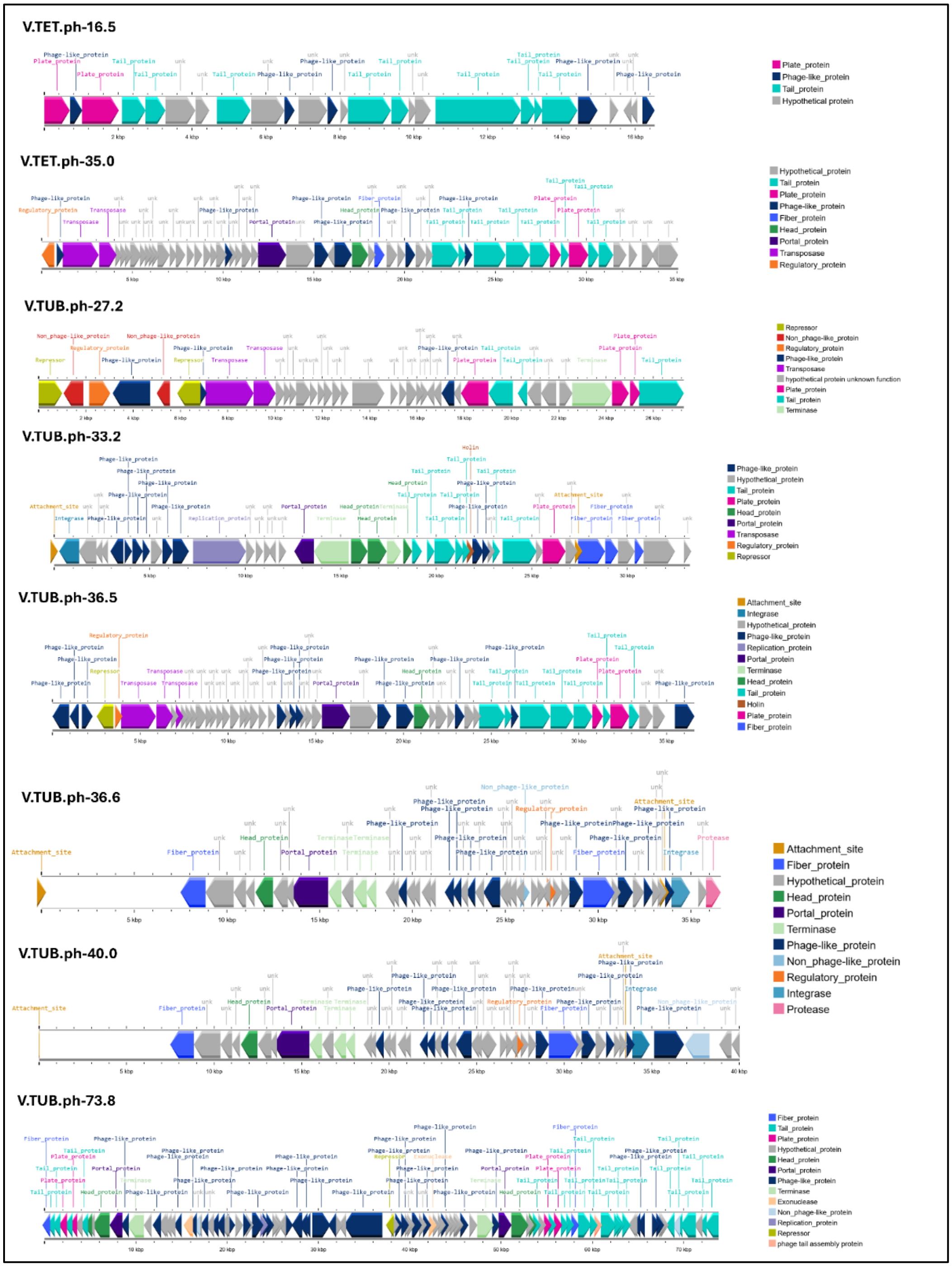


**Supplementary Figure S4. Genomic organization of intact prophages identified in *Vibrio* genomes from this study.** Gene maps of intact prophages detected using PHASTER in thirteen *Vibrio* strains isolated from *Porites cylindrica*. Prophage names correspond to those listed in Table 4. Arrows represent predicted open reading frames and indicate transcriptional orientation. Genes are color-coded by functional category (e.g., head proteins, tail proteins, portal proteins, terminase, integrase, replication proteins, regulatory proteins, transposases, hypothetical proteins). Scale bars indicate genomic distance (kb). These prophages vary in size and gene composition but share conserved structural modules typical of temperate bacteriophages. While most prophages encode structural and replication functions, only a subset carry identifiable virulence-associated genes (see Figure 7), highlighting functional heterogeneity among prophage elements.

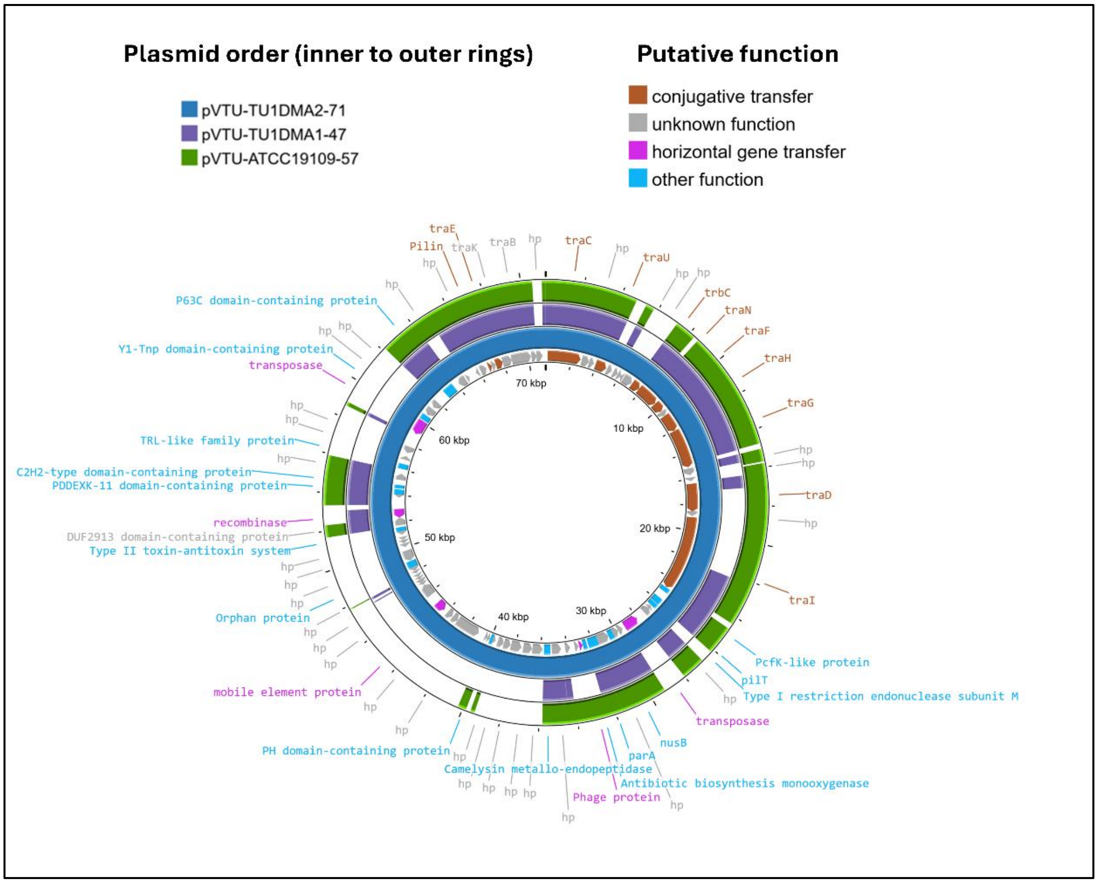


**Supplementary Figure S5**. Comparative circular alignment of plasmids from 3 *Vibrio tubiashii* strains, including two plasmids sequenced in this study (pVTU-TU1MA2-71 and pVTU-TU1MA1-47) and a reference plasmid from the NCBI database (pVTU-ATCC19109-57; see Table 3 for strain information and accession number). Each concentric ring represents a single plasmid and is labeled with its corresponding plasmid name. Colored features indicate functional gene categories, including conjugative transfer systems (brown), horizontal gene transfer–associated genes (pink), other functions (blue), and unknown functions (gray).
